# Short-course but not prolonged treatment with ATR inhibitor AZD6738 integrates with radiotherapy to generate a tumor antigen-specific CD8^+^ T cell expansion in the periphery

**DOI:** 10.1101/2022.04.11.487886

**Authors:** Frank P. Vendetti, David A. Clump, Sandra Schamus-Haynes, Maria DiMayorca, Naveed Islam, Jina Chang, Jan H. Beumer, Christopher J. Bakkenist

**Author notes:** Corresponding Authors: Frank P. Vendetti, Hillman Cancer Center, 2.1, 5117 Centre Avenue, Pittsburgh, PA 15232, Christopher J. Bakkenist, Hillman Cancer Center, 2.6, 5117 Centre Avenue, Pittsburgh, PA 15232.

## Abstract

ATR kinase is a central regulator of the DNA damage response. While ATR kinase inhibitors (ATRi’s) are known to sensitize cancer cells to DNA damage, their effect on immune cells is not known. Here we show in mice that short-course AZD6738 (ATRi) on days 1-3 decreases proliferating T cells in the tumor and periphery and that cessation of ATRi causes a proliferative rebound. Integrating radiation on days 1-2 (RT) with ATRi on days 1-3 increases IFN-β in the tumor and activates tumor antigen-specific CD8^+^ T cells in the tumor-draining lymph node (DLN). RT with short-course ATRi induces an expansion of tumor antigen-specific CD8^+^ T cells in the DLN. In contrast, RT with prolonged daily ATRi blocks expansion of antigen-specific CD8^+^ T cells, despite increased IFN-β and activation of CD8^+^ T cells. Our data identifies critical schedule considerations for ATRi with RT, immunotherapy and genotoxic therapies.

## Introduction

Most chemotherapies target DNA replication forks and their efficacy is associated with both the direct killing of proliferating tumor cells and, in many cases, the stimulation of innate and adaptive anti-tumor immune responses (*1*, *2*). One mechanism through which DNA damaging chemotherapies increase anti-tumor immune responses is the direct killing of proliferating immune cells causing transient leukocytopenia followed by a rebound proliferation of immune cell populations. A major clinical goal is to identify novel regimens that maximize the direct killing of tumor cells, while concurrently selecting for and stimulating the activity of anti-tumor immune effector cells.

The DNA damage response (DDR) is a signaling network that recognizes damaged DNA and coordinates cellular responses to control cell cycle progression, DNA repair, and cell death (*3*). Pharmacologic inhibitors of DDR kinases (DDRi) have been developed with the goal of enhancing tumor cell killing by DNA-damaging therapies (*2*, *4*). Advances in our understanding of how the DDR in tumor cells impacts immune responses has additionally generated interest in combining DDRi with immunotherapy (*5*–*7*). However, the direct effect of DDRi in immune cells has not been investigated.

ATR is a DDR kinase activated at damaged replication forks and resected DNA double-strand breaks (DSBs). Six ATR kinase inhibitors have advanced to phase 1 and phase 2 trials: ceralasertib (AZD6738); berzosertib (M6620, VX-970); elimusertib (BAY 1895344); RP-3500; M4344 (VX-803); and M1774 (*8*–*13*). In these clinical investigations, ATR kinase inhibitors are being combined with genotoxic chemotherapy, poly (ADP-ribose) polymerase (PARP) inhibitors, radiotherapy, and immunotherapy, and substantial preclinical evidence demonstrates that AZD6738 (ATRi) sensitizes cells to DNA damaging agents (*14*–*24*). More recently, preclinical studies have also shown that ATRi can potentiate anti-tumor immune responses, but the underlying mechanisms are not well-understood (*17*, *19*, *25*–*28*).

Preliminary phase I data show that ATRi suppresses circulating monocytes and proliferating T cells in patients, and that neutropenia and thrombocytopenia are dose-limiting and schedule-limiting toxicities of ATRi (*29*–*32*). Nevertheless, in several of the 25 active clinical trials of ATRi, patients treated with immunotherapy are given ATRi twice-daily for up to two weeks (*33*–*41*). Preclinical investigations of the impact of ATRi in immune cells *in vivo*, both in the tumor microenvironment and periphery, are urgently needed, as the impairment of proliferating immune cells, and particularly CD8^+^ T cells, by ATRi could hamper responses to immunotherapy.

We previously demonstrated that ATRi combines with conformal tumor radiotherapy (RT) to generate durable, CD8^+^ T cell dependent anti-tumor responses in mouse models of cancer (*17*). During this work, we observed that short-course, 3-day treatment with ATRi, independent of RT, transiently reduced proliferating T cell populations in spleens and tumor infiltrating lymphocytes (TIL) as well as activated, effector/effector memory CD8^+^ T cells in spleens of tumor-bearing mice. Here we systematically investigate the effect of short-course ATRi and prolonged ATRi given daily on T cell responses in TIL and the periphery.

## Results

### Short-course ATRi treatment reduces circulating leukocytes and decreases proliferating T cells in the tumor infiltrate and peripheral lymphoid tissues

To better understand how ATRi impacts immune populations in mice, we first treated BALB/c mice with 75 mg/kg AZD6738 on days 1-3 (ATRi QDx3) and collected peripheral blood at day 4 for complete blood counts (CBC). ATRi QDx3 treatment significantly reduced circulating total white blood cells (WBC), lymphocytes, monocytes, and neutrophils (Fig 1A). Therefore, a short-course, 3-day ATRi treatment reduces all circulating white blood cell populations examined, and this is consistent with the effects of ATRi on WBC in patients in phase I trials (*29*–*32*).

**Figure 1.**
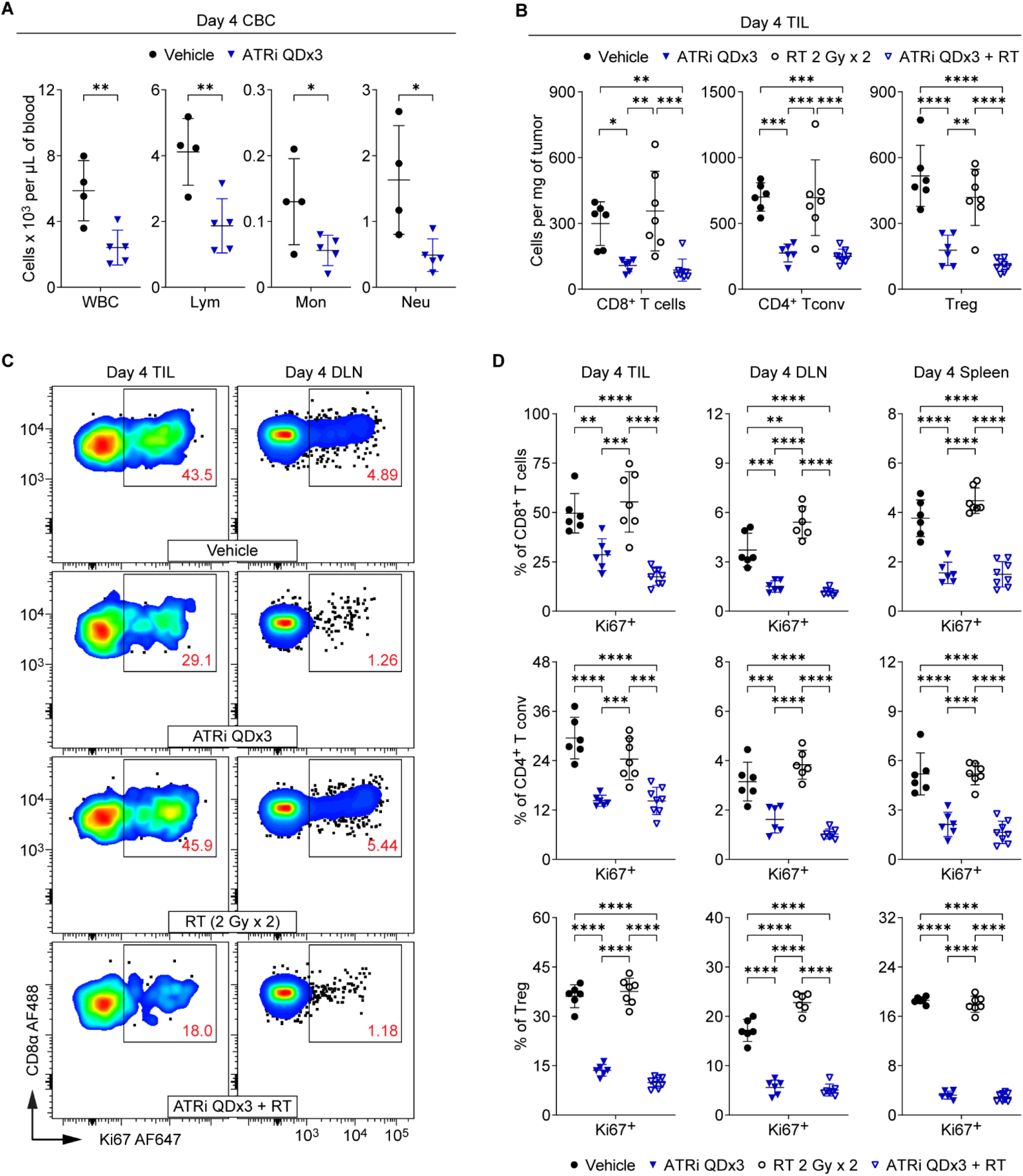
Short-course ATRi treatment systemically impacts leukocytes and depletes the tumor and peripheral lymphoid tissues of proliferating T cells. **A**. Complete blood counts (CBC) at Day 4 from BALB/c mice treated with ATRi on days 1-3 (ATRi QDx3) or vehicle. n = 4 vehicle, 5 ATRi QDx3. Mean and SD bars shown. *p<0.05, **p<0.01 by unpaired, two-tailed t test. **B-D**. CT26 tumor-bearing mice were treated with ATRi on days 1-3 (ATRi QDx3), radiotherapy on days 1-2 (RT 2 Gy x 2), ATRi QDx3 + RT, or vehicle. **B**. Quantitation of the number of CD8^+^ T cells, conventional CD4^+^ T cells (CD4^+^ Tconv), and regulatory T cells (Treg) in the tumor-infiltrating leukocytes (TIL), per mg of tumor, at Day 4. **C**. Representative cytograms depicting expression of Ki67 in CD8^+^ T cells in the TIL and tumor-draining lymph node (DLN) at Day 4**. D**. Quantitation of proliferating (Ki67^+^) CD8^+^ T cells, CD4^+^ Tconv, and Treg, as a percentage of the corresponding parent population, in the TIL, DLN, and Spleen at Day 4. **B,D**. Data from 3 independent experiments with 1-3 mice per group. n = 6 Vehicle, 6 ATRi QDx3, 7 RT (6 DLN), 8 ATRi QDx3 + RT (7 DLN). Mean and SD bars shown. **p<0.01, ***p<0.001, ****p<0.0001 by ANOVA with Tukey’s multiple comparisons test.

Next, we sought to examine the impact of short-course ATRi treatment, alone or in combination with conformal tumor radiotherapy on days 1-2 (RT, 2 Gy x 2) on T cell populations in the TIL and peripheral lymphoid tissues, including spleen and tumor-draining lymph node (DLN), of CT26 tumor-bearing mice at day 4. We employed our previously established ATRi QDx3 and RT treatment protocol (*17*) and used multi-parameter flow cytometry to immunoprofile T cell populations. First, we enumerated CD8^+^ T cells, CD4^+^ conventional T cells (CD4^+^ Tconv), and regulatory T cells (Treg). ATRi QDx3 significantly reduced all three T cell populations in the TIL compared to both vehicle and RT alone (Fig 1B). This effect was independent of RT. Conversely, ATRi QDx3 did not reduce the relative number of these T cells in the peripheral lymphoid tissues (Fig S1A,B).

Next, we examined the impact of short-course ATRi treatment on proliferating T cells, using the proliferation marker Ki67. Independent of RT, ATRi QDx3 significantly reduced proliferating (Ki67^+^) CD8^+^ T cells, relative to both vehicle and RT alone, in the TIL, DLN, and spleens at day 4 (Fig 1C,D). We observed similar effects of ATRi QDx3 on proliferating CD4^+^ Tconv and Treg in all tissues examined at day 4, with a striking decrease in proliferating Tregs (Fig 1D).

Finally, we examined the impact of short-course ATRi treatment on the effector/effector memory (TEM, CD44^hi^CD62L^lo^), central memory (TCM, CD62L^hi^CD44^hi^), and naïve (TN, CD62L^hi^CD44^lo^) CD8^+^ T cell subsets, as a percentage of the total CD8^+^ T cell pool, in the DLN and spleens. In the DLN, ATRi QDx3 plus RT significantly decreased the proportion of TEM when compared to RT alone, significantly decreased the proportion of TCM when compared to vehicle, and significantly increased the proportion of TN when compared to vehicle and ATRi QDx3 alone (Fig S1C). In spleens, ATRi QDx3 plus RT significantly reduced the relative TEM subset when compared to RT alone and significantly reduced the relative TCM subset compared to all other treatments. Accordingly, ATRi QDx3 plus RT increased the relative TN subset when compared to vehicle and RT alone (Fig S1D). Collectively, these results demonstrate that short-course, 3-day ATRi treatment reduces activated CD8^+^ T cell subsets in the periphery at day 4 and are consistent with our previous findings in spleens at day 5 (*17*).

### Cessation of ATRi treatment results in a rapid proliferative rebound in T cells

Our previous work demonstrated that a rebound in proliferating (Ki67^+^) T cells occurs after cessation of short-course, 3-day ATRi treatment at some time between day 5 and day 9 (*17*). To better understand when T cells recover from ATRi treatment, and when this proliferative rebound occurs, we immunoprofiled T cell populations at day 7 and day 9. Evidence of proliferative rebound was apparent by day 7 for all T cell populations in all tissues examined (Fig 2A). CD8^+^ T cells and CD4^+^ T conv exhibited similar rebound patterns across tissues at this timepoint. Proliferating CD8^+^ T cells and proliferating CD4^+^ Tconv in the TIL of ATRi QDx3-treated and ATRi QDx3 plus RT-treated mice recovered to near or above the percentages observed in vehicle-treated mice (Fig 2A). In the DLN, these proliferating T cell populations in ATRi QDx3-treated and ATRi QDx3 plus RT-treated mice rebounded to percentages significantly higher than observed in vehicle-treated mice, and in the case of ATRi QDx3 plus RT-treated mice, significantly higher than in mice treated with RT alone (Fig 2A). In spleens, only in mice treated with ATRi QDx3 plus RT did the percentages of proliferating CD8^+^ T cells and proliferating CD4^+^ Tconv increase significantly compared to vehicle-treated mice (Fig 2A). Treg exhibited a stronger proliferative rebound at day 7, as the percentages of proliferating Treg in ATRi QDx3-treated and ATRi QDx3 plus RT-treated mice increased significantly compared to vehicle-treated mice in all tissues examined (Fig 2A).

**Figure 2.**
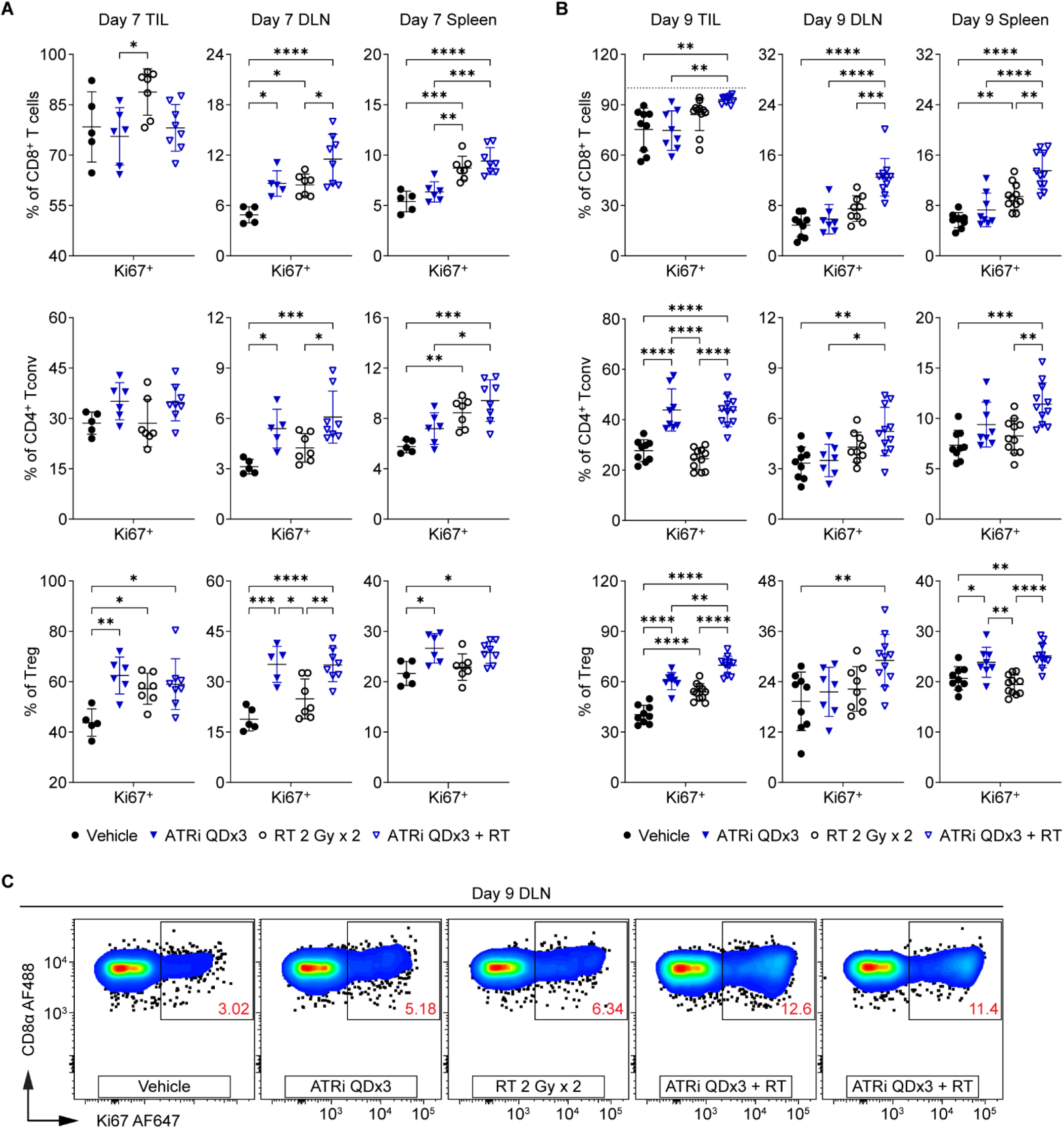
Cessation of ATRi treatment results in a rapid proliferative rebound in T cells. **A-C**. CT26 tumor-bearing mice were treated with ATRi on days 1-3 (ATRi QDx3), radiotherapy on days 1-2 (RT 2 Gy x 2), ATRi QDx3 + RT, or vehicle. **A**. Quantitation of proliferating (Ki67^+^) CD8^+^ T cells, CD4^+^ Tconv, and Treg, as a percentage of the corresponding parent population, in the TIL, DLN, and Spleen at Day 7. Data from at least 2 independent experiments with 1-4 mice per group. n = 5 Vehicle, 6 ATRi QDx3 (5 DLN), 7 RT, 8 ATRi QDx3 + RT. **B**. Quantitation of proliferating (Ki67^+^) CD8^+^ T cells, CD4^+^ Tconv, and Treg, as a percentage of the corresponding parent population, in the TIL, DLN, and Spleen at Day 9. Data from at least 4 independent experiments with 1-3 mice per group. n = 9 Vehicle, 8 ATRi QDx3 (7 DLN), 11 RT (9 DLN), 11 ATRi QDx3 + RT. **A-B**. Mean and SD bars shown. *p<0.05, **p<0.01, ***p<0.001, ****p<0.0001 by ANOVA with Tukey’s multiple comparisons test. **C**. Representative cytograms depicting expression of Ki67 in CD8^+^ T cells in DLN at day 9. Two examples, representing two independent mice, are shown for ATRi QDx3 + RT.

While proliferative rebound in all T cell populations was apparent by day 7, none of these T cell populations in the TIL of ATRi QDx3-treated and ATRi QDx3 plus RT-treated mice had fully recovered to vehicle control numbers (Fig S2A). This was most evident with the CD4^+^ Tconv and Treg populations in ATRi QDx3 plus RT-treated treated mice, which were significantly reduced compared to vehicle-treated and RT-treated mice. In the periphery, relative Treg numbers were comparable across all treatments. CD8^+^ T cells and CD4^+^ Tconv were reduced in the DLN by ATRi QDx3, with or without RT, compared to vehicle, but were unchanged in spleens (Fig S2B). As our reported quantitation of T cell populations is relative, and not absolute, the differences observed between DLN and spleen are likely the result of the different immune cell composition of these tissues and any differential effects of ATRi, or in the timing of recovery from ATRi, among the different immune cell types.

At day 9, all T cell populations in all examined tissues from ATRi QDx3 plus RT-treated mice exhibited significant increases in the percentages of proliferating T cells compared to vehicle-treated mice (Fig 2B). ATRi QDx3 alone increased the percentages of proliferating CD4^+^ Tconv and Treg in the TIL, and proliferating Treg in spleens, but did not increase proliferating CD8^+^ T cells in any tissues (Fig 2B). More striking was that ATRi QDx3 plus RT-treated mice exhibited significant increases in proliferating CD8^+^ T cells in DLN and spleens compared to RT-treated mice (Fig 2B,C). Therefore, our findings at day 9 indicate a persistent proliferative response in CD8^+^ T cells in the periphery that requires both RT and ATRi QDx3.

Examination of T cell numbers at day 9 revealed evidence of further repopulation of the TIL in ATRi QDx3 plus IR-treated mice, and the number of CD8^+^ T cells recovered to at least control levels. (Fig S3A). The relative numbers of T cells in the peripheral tissues had largely normalized, and we found no significant differences in CD8^+^ T cells in DLN or spleens for any treatment (Fig S3B,C). Collectively, these data demonstrate that a proliferative rebound occurs within 4 days of cessation of short-course ATRi treatment. For CD4^+^ Tconv and Treg, increases in proliferating cells persist in some tissues to day 9. However, for CD8^+^ T cells, there is no evidence of a continued proliferative rebound at day 9 following ATRi QDx3 alone, and the proliferative response that persists in the TIL, DLN, and spleens requires both RT and ATRi QDx3.

### Short-course ATRi integrates with radiotherapy to promote expansion of tumor antigen specific CD8^+^ T cells in the tumor-draining lymph node

We next examined whether the persistent proliferation of CD8^+^ T cells in the periphery at day 9 after ATRi QDx3 plus RT was associated with an increase in activated, effector CD8^+^ T cells in the periphery. ATRi QDx3 plus RT significantly increased the proportion of TEM in the DLN compared to all other treatments (Fig 3A,B). This correlated with a significant reduction in the proportion of TN in the DLN of ATRi QDx3 plus RT-treated mice compared to vehicle-treated mice (Fig 3B). In spleens, ATRi QDx3 plus RT significantly increased the TEM subset compared to vehicle and ATRi QDx3 alone, but not compared to RT alone, which also increased the TEM pool compared to vehicle control (Figure S4A). No significant differences among treatments were observed in the proportions of TCM in the DLN or spleens, or in the proportions of TN in spleens (Fig 3B,S4A). Thus, expansion of an effector/effector memory (TEM) CD8^+^ T cell pool at day 9 in the DLN occurs only in ATRi QDx3 plus RT-treated mice.

**Figure 3.**
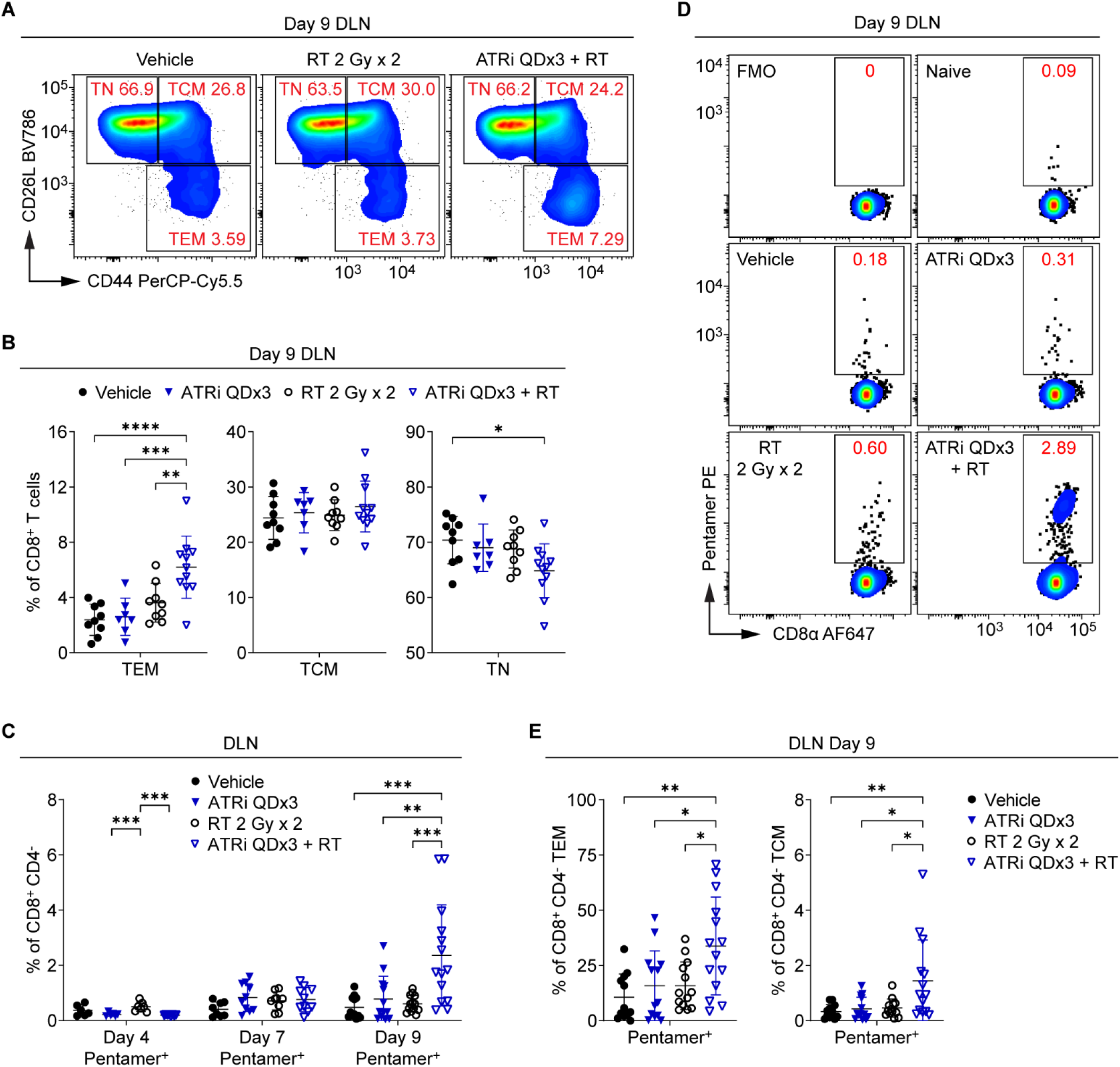
Short-course ATRi integrated with radiotherapy promotes expansion of tumor antigen specific CD8^+^ T cells in the tumor-draining lymph node. **A-E**. CT26 tumor-bearing mice were treated with ATRi on days 1-3 (ATRi QDx3), radiotherapy on days 1-2 (RT 2 Gy x 2), ATRi QDx3 + RT, or vehicle. **A**. Representative cytograms depicting expression of CD62L and CD44 on CD8^+^ T cells in the DLN at day 9. CD8^+^ T cell subpopulations were defined as effector/effector memory (TEM, CD44^hi^CD62L^lo^), central memory (TCM, CD62L^hi^CD44^hi^), or naïve (TN, CD62L^hi^CD44^lo^). **B**. Quantitation of CD8^+^ TEM, TCM, and TN, as a percentage of total CD8^+^ T cells, in the DLN at day 9. Data from at least 4 independent experiments with 1-3 mice per group. n = 9 Vehicle, 7 ATRi QDx3, 9 RT, 11 ATRi QDx3 + RT. **C**. Quantitation of tumor antigen specific (AH1 Pentamer^+^) CD8^+^ T cells, as a percentage of total CD8^+^CD4^-^ cells, in the DLN at Day 4, Day 7, and Day 9. Data from 3 (Day 4), at least 3 (Day 7), and at least 5 (Day 9) independent experiments with 1-5 mice per group. n at Day 4 = 6 Vehicle, 6 ATRI QDx3, 7 RT, 8 ATRi QDx3 + RT. n at Day 7 = 8 Vehicle, 10 ATRi QDx3, 10 RT, 11 ATRi QDx3 + RT. n at Day 9 = 12 Vehicle, 13 ATRi QDx3, 13 RT, 14 ATRi QDx3 + RT. **D**. Representative cytograms depicting AH1 Pentamer^+^ CD8^+^ T cells in the DLN at day 9. Fluorescence-minus-one (no Pentamer PE) and Naïve (negative control, no tumor) controls also shown. **E**. Quantitation of AH1 Pentamer^+^ CD8^+^ TEM and TCM, as a percentage of total CD8^+^CD4^-^ TEM and TCM, in the DLN at day 9. **B,C,E**. Mean and SD bars shown. *p<0.05, **p<0.01, ***p<0.001, ****p<0.0001 by ANOVA with Tukey’s multiple comparisons test.

Next, we sought to determine whether expansion of CD8^+^ TEM cells in the DLN at day 9 was nonspecific or was associated with expansion of tumor antigen-specific CD8^+^ T cells. To identify CD8^+^ T cells specific for CT26 tumor antigen, we utilized a PE-labeled, pentameric MHC class I H-2Ld complex loaded with the AH1 peptide (SPSYVYHQF). This peptide sequence is specific for the murine leukemia virus (MuLV) gp70 envelope protein, which is the immunodominant antigen expressed by CT26 (*42*–*44*). We enumerated antigen-specific CD8^+^ T cells, as a percentage of the total CD8^+^CD4^-^ cell pool, in the DLN at day 4, day 7, and day 9 following treatment with ATRi QDx3, RT, ATRi QDx3 plus RT, or vehicle. At day 4, the overall percentages pentamer^+^ CD8^+^ T cells were low among all treatment groups. ATRi QDx3, alone or plus RT, further reduced pentamer^+^ CD8^+^ T cells compared to RT alone (Fig3C). This occurred despite a relative increase in total CD8^+^CD4^-^ cells, as a percentage of all CD45^+^ immune cells, after ATRi QDx3 and ATRi QDx3 plus IR compared to IR alone (Fig S4B).

We found no significant differences among treatments in the percentages of pentamer^+^ CD8^+^ T cells in the DLN at day 7, whereas at day 9, pentamer^+^ CD8^+^ T cells were significantly increased in the DLN of ATRi QDx3 plus RT-treated mice compared to all other treatment groups (Fig 3C,D). This increase occurred absent any significant change in the relative amount of total CD8^+^CD4^-^ cells in the DLN (Fig S4B). In addition, our naïve (no tumor) mouse negative controls, included with each experiment, demonstrated very low non-specific binding (Fig 3D). Therefore, ATRi QDx3 plus RT promotes expansion of tumor antigen-specific CD8^+^ T cells in the DLN by day 9.

Next, we questioned whether the expanding tumor antigen-specific CD8^+^ T cells possess a TCM or TEM phenotype at this timepoint. ATRi QDx3 plus RT significantly increased both the pentamer^+^ CD8^+^CD4^-^ TEM and pentamer^+^ CD8^+^CD4^-^ TCM populations compared to all other treatment groups (Fig 3E). Antigen-specific CD8^+^ TEM comprised a large portion of the total CD8^+^CD4^-^ TEM pool, and in some ATRi QDx3 plus RT-treated mice, accounted for more than half of the total effector/effector memory CD8^+^ T cell pool (Fig 3E). Therefore, short-course ATRi plus RT promotes expansion of tumor antigen-specific, effector/effector memory CD8^+^ T cells in the DLN by day 9.

### Short-course ATRi treatment potentiates radiotherapy-induced type I interferon signaling and inflammatory cytokines and chemokines in the tumor microenvironment

Our previous work using this CT26 model demonstrated that ATRi QDx3 plus RT results in more functional CD8^+^ T cells in the TIL at later timepoints (*17*). We now demonstrate ATRi QDx3 plus RT promotes the expansion of tumor antigen-specific CD8^+^ T cell in the DLN at day 9. We sought to better understand what drives this adaptive CD8^+^ T cell response in the periphery following ATRi QDx3 plus RT. Recent preclincal studies show that ATRi increases type I interferon (IFN) signaling and production of proinflammatory cytokines and chemokines after radiation *in vitro* and *in vivo (19, 26*). Similar proinflammatory signaling occurs after DNA damage induced by combined ATR and WEE1 inhibition (*28*). Furthermore, combined ATRi and radiotherapy is associated with increased antigen presentation on tumor cells and myeloid immune cell infiltration of tumors *in vivo (19*). Collectively, the preclinical data suggest that proinflammatory cytokine signaling and innate immune system activity following ATRi plus RT-induced DNA damage are linked to subsequent priming of the adaptive anti-tumor immune response.

To examine type I IFN signaling and production of proinflammatory cytokines and chemokines in the tumor microenvinroment, we employed a customized 10-plex immunoassay on the U-Plex platform to measure levels of the following 10 cytokines/chemokines in tumor extract: GM-CSF, IFN-α, IFN-β, IFN-γ, IL-6, IL-12p70, CXCL10 (IP-10), CCL2 (MCP-1), CCL3 (MIP-1α), and CCL5 (RANTES). These targets were selected for several reasons. First, we wanted to examine both type I (IFN-α and IFN-β) and type II IFN (IFN-γ) signaling in the tumor microenvironment. Second, expression of IFN-β, the proinflammatory cytokine IL-6, and the proinflammatory chemokines CXCL10, CCL2, and CCL5 were previously reported to be strongly induced in tumor cells by ATRi plus RT compared to RT alone (*19*, *26*). In addition, ATRi plus RT enhanced secretion of CXCL10, CCL3, and CCL5 by tumor cells *in vitro* (*19*). We also included the immune modulatory cytokine GM-CSF, which is secreted by a variety of cells, including immune cells and cancer cells, due to its functions in regulating the maturation and functional activity of monocytes, macrophages, and dendritic cells (*45*, *46*). Lastly, we included the proinflammatory cytokine IL-12p70, which is produced primarily by dendritic cells and macrophages and promotes NK and T cell-mediated IFN-γ production and cytotoxicity (*47*).

We treated mice with RT, ATRi QDx3 plus RT, or vehicle, and measured levels of cytokines/chemokines in extracts from tumors harvested at day 5 (Fig 4A) and day 7 (Fig 4B). The ATRi QDx3 alone treatment group was omitted from these studies as our data showed no increase in peripheral CD8^+^ TEM expansion or peripheral tumor antigen-specific CD8^+^ T cell expansion with ATRi QDx3 alone. In addition, data from other models support that RT and ATRi are both required for the full inflammatory response (*19*, *26*). At day 5, RT alone and ATRi QDx3 plus RT significantly increased levels of GM-CSF, CXCL10, CCL2, and CCL3 compared to vehicle, but addition of ATRi QDx3 did not potentiate levels of these proteins beyond RT alone (Fig 4C and S5A). RT alone also singificantly increased levels if IFN-γ compared to vehicle and ATRi QDx3 plus RT (Fig 4C). ATRi QDx3 attenuated RT-induced IFN-γ in the tumor microenvironment, consistent with our prior findings that ATRi QDx3 attenutated both RT-induced PD-L1 expression on tumor cells and RT-induced increases in IFN-γ competent CD8^+^ T cells in the TIL at this timepoint (*17*). At day 5, ATRi QDx3 plus RT significantly increased levels of IFN-β and CCL5 compared to vehicle control, but not compared to RT (Fig 4C). No changes in IFN-α, IL-6, or IL-12p70 were observed at day 5 (Fig S5A).

**Figure 4.**
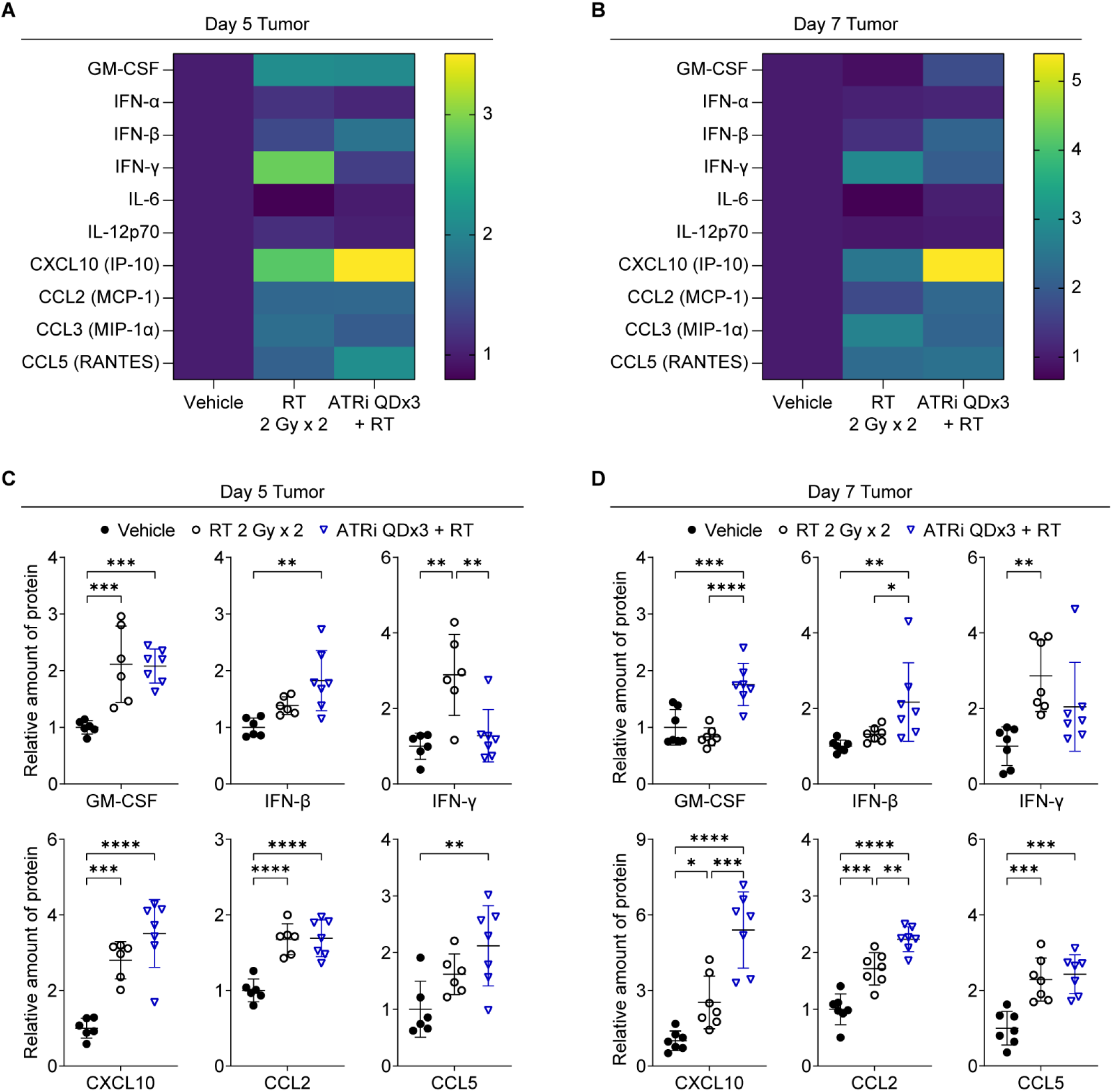
Short-course ATRi treatment potentiates radiotherapy-induced type I interferon signaling and inflammatory cytokines and chemokines in the tumor microenvironment. **A-D**. CT26 tumor-bearing mice were treated with RT (2 Gy x 2), ATRi QDx3 + RT, or vehicle. **A**. Heatmaps depicting the mean relative expression (compared to vehicle control) of 10 inflammatory cytokines and chemokines in tumors at Day 5. **B**. Heatmaps depicting mean relative expression (compared to vehicle control) of 10 inflammatory cytokines and chemokines in tumors at Day 7. **C**. Quantitation of the relative amount of protein (normalized to vehicle control) of a subset of inflammatory cytokines and chemokines in tumors at Day 5 **D**. Quantitation of the relative amount of protein (normalized to vehicle control) of a subset of inflammatory cytokines and chemokines in tumors at Day 7. **A,C**. Data from one experiment. n = 6 Vehicle, 6 RT, 7 ATRi QDx3 + RT. **B,D**. Data from two independent experiments with 3-4 mice per group. n = 7 Vehicle, 7 RT, 7 ATRi QDx3 + RT. **C,D**. Mean and SD bars shown. *p<0.05, **p<0.01, ***p<0.001, ****p<0.0001 by ANOVA with Tukey’s multiple comparisons test.

At day 7, ATRi QDx3 plus RT significantly increased GM-CSF and IFN-β relative to both RT alone and vehicle control and potentiated RT-induced increases in CXCL10, and CCL2 (Fig 4D). CCL5 was significantly increased by both RT alone and ATRi QDx3 plus RT, but ATRi QDx3 did not potentiante levels of CCL5 over RT alone (Fig D). ATRi QDx3 plus RT did not alter levels of IFN-α, IL-6, IL-12p70, or CCL3 at day 7 (Fig S5B). Therefore, short-course ATRi plus RT increases type I IFN (IFN-β) and several proinflammatory cytokines/chemokines, including GM-CSF, CXCL10, and CCL2 in the tumor microenvironment, and these results are consistent with published findings in other model systems (*19*, *26*).

### Short-course ATRi plus radiotherapy promotes accumulation of inflammation-associated innate immune cells and increased CD8^+^ T cell activation in the DLN

Next we sought to determine whether increased proinflammatory signaling in the tumor microenvironment was associated with changes in the DLN that may explain the later expansion of tumor antigen-specific CD8^+^ T cells in the DLN. Therefore, we immunoprofiled DLN from the same mice whose tumors were harvested for cytokine and chemokine measurements at the day 7 timepoint. First, we examined for evidence of altered migration of innate immune cells to the DLN after ATRi QDx3 plus RT. After gating out all T and B lymphocytes, CD3^-^CD19^-^ cells were profiled to quantify NK cells (NKp46^+^), monocytes and macrophages (mon/macro, CD11b^+^ and NKp46^-^CD11c^-^), and total dendritic cells (DC, CD11c^+^ and NKp46^-^). ATRi QDx3 plus RT significantly increased NK cells in the DLN, but did not change the the mon/macro, DC, and overall CD3^-^CD19^-^ cell populations (Fig 5A and S5C).

**Figure 5.**
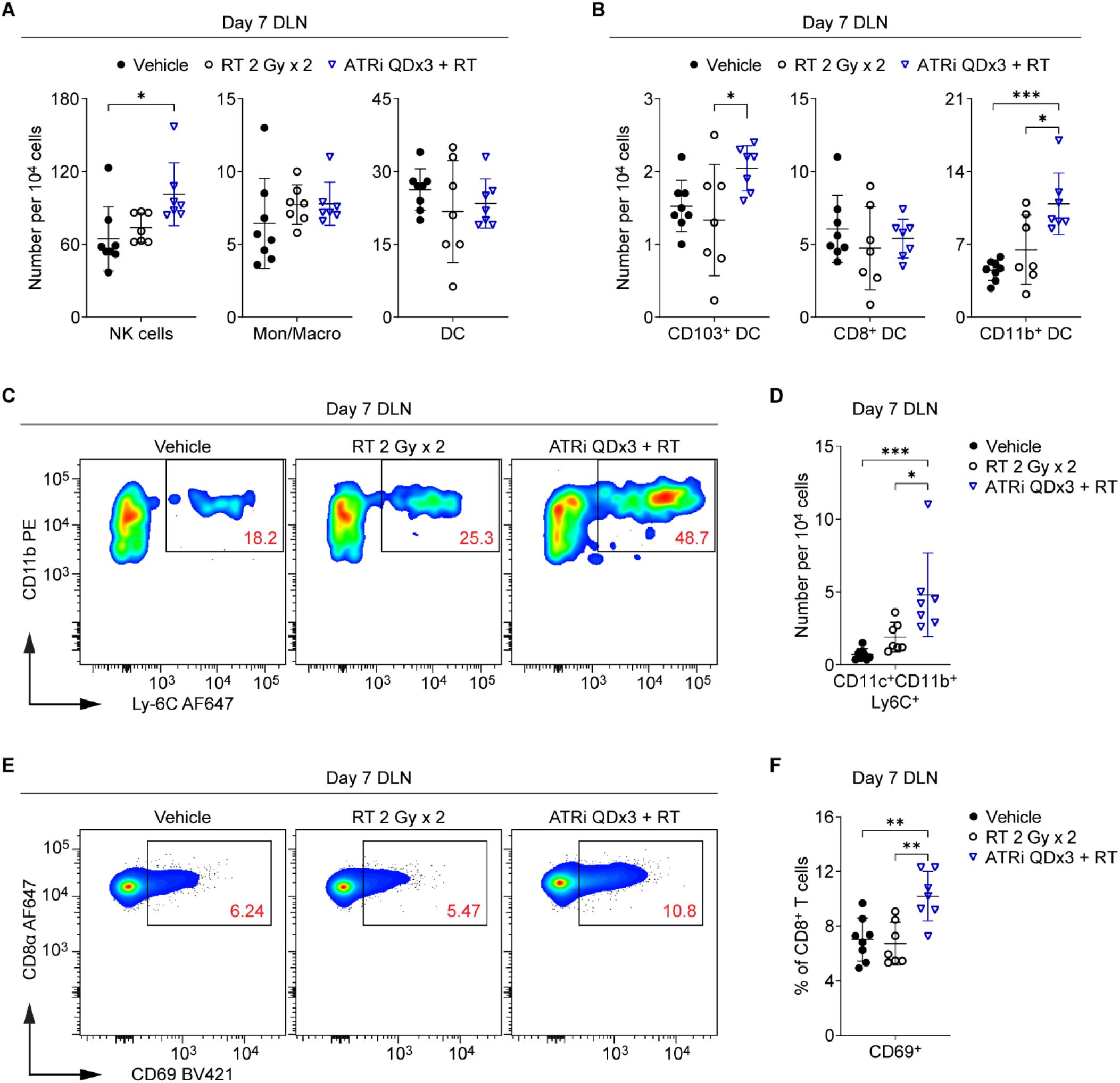
Short-course ATRi plus radiotherapy promotes accumulation of inflammation-associated innate immune cells and increased CD8^+^ T cell activation in the DLN. **A-F**. CT26 tumor-bearing mice were treated with RT (2 Gy x 2), ATRi QDx3 + RT, or vehicle. **A**. Quantitation of the relative numbers of innate immune cell populations in the DLN at Day 7. Immunoprofiled populations include NK cells (NKp46^+^ and CD3^-^CD19^-^), monocytes/macrophages (mon/macro, CD11b^+^ and CD3^-^CD19^-^NKp46^-^CD11c^-^), and dendritic cells (DC, CD11c^+^ and CD3^-^CD19^-^ NKp46^-^). **B**. Quantitation of the relative numbers of dendritic cell (DC) subsets in the DLN at Day 7. Immunoprofiled DC subsets include CD103^+^ DC (CD11c^+^CD103^+^ and CD3^-^CD19^-^NKp46^-^), CD8^+^ DC (CD11c^+^CD8^+^ and CD3^-^CD19^-^NKp46^-^CD11b^-^), and CD11b^+^ DC (CD11c^+^CD11b^+^ and CD3^-^CD19^-^ NKp46^-^CD8^-^). **C**. Representative cytograms depicting CD11b and Ly-6C expression on CD11c^+^ (and CD3^-^CD19^-^NKp46^-^) cells in the DLN at Day 7. **D**. Quantitation of the relative numbers of CD11c^+^CD11b^+^Ly-6C^+^ (and CD3^-^CD19^-^NKp46^-^CD8^-^) cells in the DLN at Day 7. **E**. Representative cytograms depicting CD69 expression on CD8^+^ T cells in the DLN at Day 7. **F**. Quantitation of newly or recently activated CD69^+^ CD8^+^ T cells, as a percentage of total CD8^+^ T cells, in the DLN at Day 7. **A,B,D,F**. Data from two independent experiments with 3-4 mice per group. n = 8 Vehicle, 7 RT, 7 ATRi QDx3 + RT. Mean and SD bars shown. *p<0.05, **p<0.01, ***p<0.001 by ANOVA with Tukey’s multiple comparisons test.

We further examined subpopulations of DC within the overal DC pool. While ATRi QDx3 plus RT did not alter the relative number of CD103^+^ DC or CD8^+^ DC (CD11c^+^CD8^+^CD11b^-^), ATRi QDx3 plus RT significantly increased the relative number of CD11b^+^ DC (CD11c^+^CD11b^+^CD8^-^) compared to both vehicle and RT (Fig 5B). We examined expression of Ly-6C within the CD11c^+^CD11b^+^ population (Fig 5C). Ly-6C expressing CD11c^+^CD11b^+^ cells are monocyte-lineage cells that arise at sites of inflammation, and cells expressing these markers fall along the differentiation specturm between inflammatory monocytes and monocyte-derived dendritic cells with functional antigen-presenting ability (*48*–*57*). ATRi QDx3 plus RT significantly increased the relative number of CD11c^+^CD11b^+^Ly-6C^+^ cells in the DLN at day 7 compared to both vehicle and RT (Fig 5C,D). These data suggest that ATRi QDx3 plus RT promotes migration of inflammation-associated innate immune cells to the DLN.

We also immunoprofiled CD8^+^ T cells to examine for evidence of T cell priming and activation in the DLN at day 7, using expression of the early activation marker CD69 on CD8^+^ T cells to identify newly or recently activated CD8^+^ T cells (Fig 5E) (*58*, *59*). Despite a reduced overall number of CD8^+^ T cells in the DLN of ATRi QDx3 plus RT-treated mice compared to vehicle control (Fig S5D), ATRi QDx3 plus RT significantly increased the percentage of the CD69^+^ CD8^+^ T cells in the DLN at day 7 compared to vehicle and RT alone (Fig 5E,F). Collectively, these data suggest that ATRi QDx3 plus RT promotes priming and activation of CD8^+^ T cells, concomitant with increased accumulation of inflammation-associated innate immune cells, in the DLN.

### Prolonged daily ATRi potentiates RT-induced IFN-β but not inflammatory chemokines in the tumor microenvironment and promotes CD8^+^ T cell activation in the DLN

Our data demonstrate that cessation of short-course, 3-day ATRi treatment triggers a proliferative rebound in T cell populations systemically, and, in parallel, ATRi QDx3 integrates with RT to increase inflammatory cytokine/chemokine signaling in the tumor microenvironment. Therefore, we sought to determine how prolonged daily ATRi treatment impacts both of these phenoma, and ultimately, how it affects the generation of the tumor antigen-specific CD8^+^ T cell response in the periphery. We first examined the impact of ATRi given for 7 consecutive days (ATRi QDx7) on RT-induced proinflammatory signalling in the tumor microenviroment at day 7 (Fig 6A). Here we also included an ATRi QDx7 alone treatment group to determine if prolonged daily ATRi in the absence of RT can trigger inflammatory signaling in tumors. Similar to short-course ATRi treatment, ATRi QDx7 plus RT significantly increased IFN-β in the tumor microenvironment compared to all other treatments, attenuated RT-induced IFN-γ, preserved IL-6 levels at baseline, and did not alter IFN-α (Fig 6B and S6). In contrast to short-course ATRi treatment, ATRi QDx7 plus RT did not potentiate RT-induced increases in CXCL10 and CCL2 (Fig 6B). In addition, ATRi QDx7 alone significantly reduced CCL5 compared to all other treatments, and ATRi QDx7 attenuated RT-induced CCL3 expression (Fig 6B).

**Figure 6.**
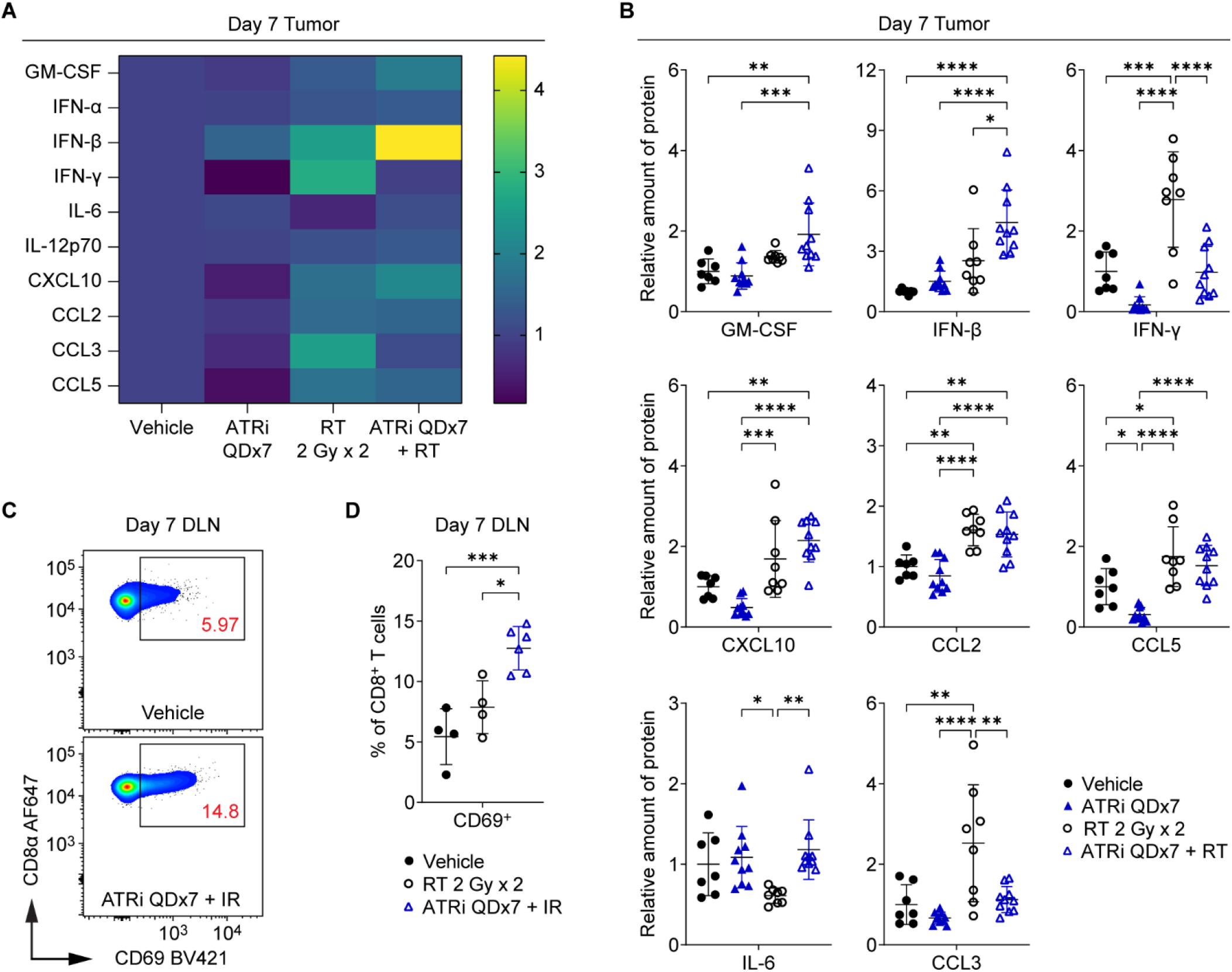
Prolonged daily ATRi potentiates RT-induced IFN-β but not inflammatory chemokines in the tumor microenvironment and promotes CD8+ T cell activation in the DLN. **A-D**. CT26 tumor-bearing mice were treated with ATRi on days 1-7 (ATRi QDx7), radiotherapy on days 1-2 (RT 2 Gy x 2), ATRi QDx7 + RT, or vehicle. **A**. Heatmaps depicting the mean relative expression (compared to vehicle control) of 10 inflammatory cytokines and chemokines in tumors at Day 7. **B**. Quantitation of the relative expression (compared to vehicle control) of a subset of inflammatory cytokines and chemokines in tumors at Day 7. **A-B**. Data from at least 4 independent experiments with 1-4 mice per group. n = 7 Vehicle, 10 ATRi QDx7, 8 RT, 10 ATRi QDx7 + RT. **C-D**. CT26 tumor-bearing mice were treated with RT (2 Gy x 2), ATRi QDx7 + RT, or vehicle. **C**. Representative cytograms depicting CD69 expression on CD8^+^ T cells in the DLN at Day 7. **D**. Quantitation of newly or recently activated CD69^+^ CD8^+^ T cells in the DLN at Day 7. Data from at least 2 independent experiments with 1-3 mice per group. n = 4 Vehicle, 4 RT, 6 ATRi QDx7 + RT. **B,D**. Mean and SD bars shown. *p<0.05, **p<0.01, ***p<0.001, ****p<0.0001 by ANOVA with Tukey’s multiple comparisons test.

We also examined CD69 expression on CD8^+^ T cells in the DLN of a subset of mice treated with RT, ATRi QDx7 plus RT, or vehicle, whose tumors were harvested for cytokine/chemokine measurements at day 7 (Fig 6C). ATRi QDx7 plus RT significantly increased the percentage of the CD69^+^ CD8^+^ T cells in the DLN at day 7 compared to vehicle and RT alone (Fig 6C,D). Therefore, ATRi QDx7 plus RT increases IFN-β signalling in the tumor microenvinroment at day 7, and despite ATRi given for 7 consecutive days, promotes early activation of CD8^+^ T cells in the DLN.

### Prolonged daily ATRi restrains the adaptive T cell response and abolishes expansion of tumor antigen-specific CD8^+^ T cells in the periphery following radiotherapy

We sought to determine the consequences of prolonged ATRi given daily on T cell infiltration of tumors. We treated mice with ATRi for 9 consectutive days (ATRi QDx9), with or without RT (2 Gy x 2), and harvested tumors on day 9. ATRi QDx9 treatment, alone or with RT, resulted in TIL that were essentially devoid of CD8^+^ T cells and Treg (Fig 7A). Similarly, the TIL of these mice had significantly reduced numbers of CD4^+^ Tconv compared to vehicle-treated and RT-treated mice (Fig 7A). What little of each T cell population that was present in the TIL of ATRi QDx9-treated and ATRi QDx9 plus RT-treated mice exhibited significantly reduced proportions of proliferating T cells (Fig 7B). Therefore, prolonged, 9-day ATRi treatment results in a dearth of proliferating and total T cells in the TIL at day 9.

**Figure 7.**
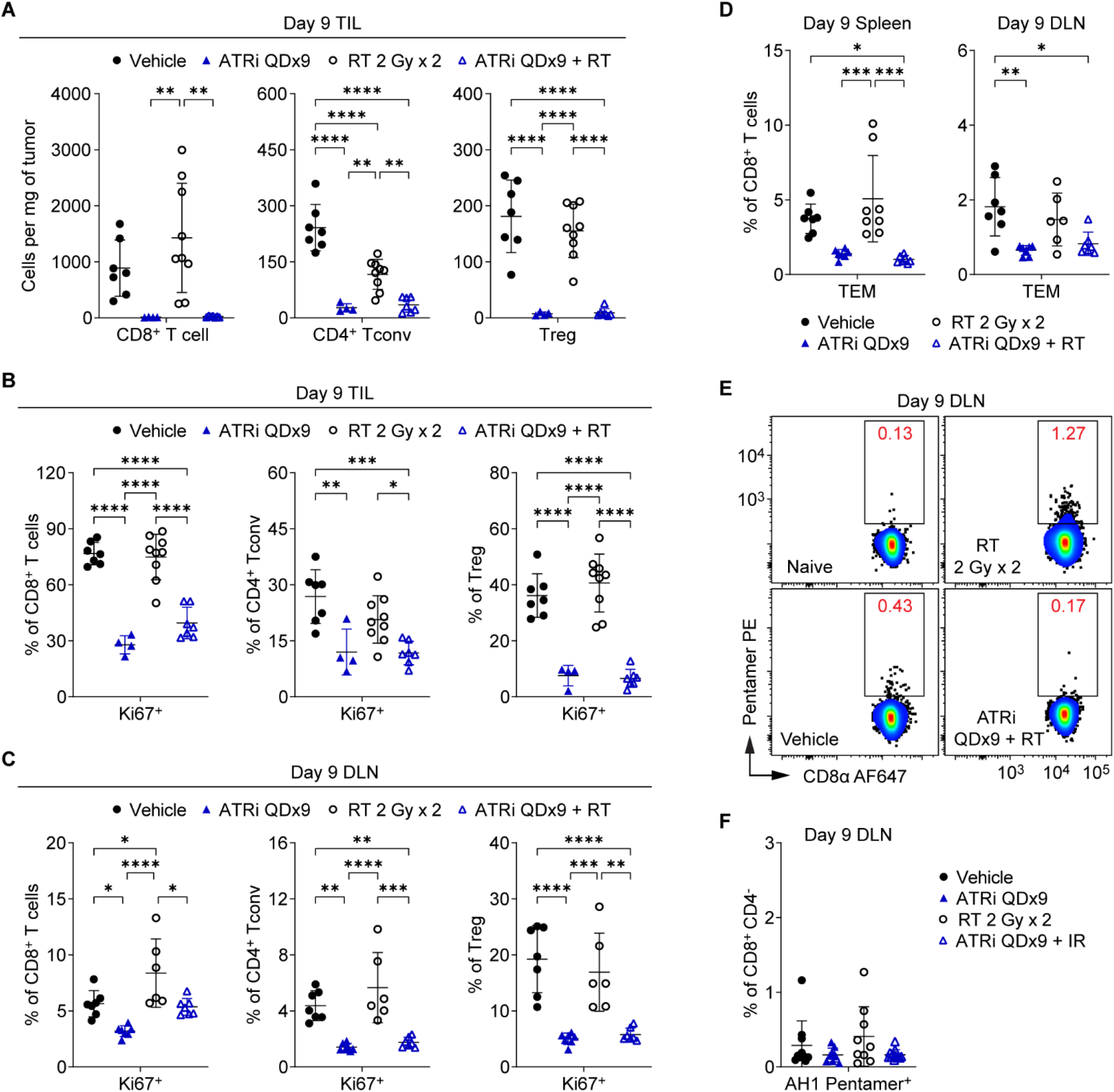
Prolonged daily ATRi treatment restrains the adaptive T cell response and abolishes expansion of tumor antigen-specific CD8^+^ T cells in the periphery following radiotherapy. **A-D**. CT26 tumor-bearing mice were treated with ATRi on days 1-9 (ATRi QDx9), radiotherapy on days 1-2 (RT 2 Gy x 2), ATRi QDx9 + RT, or vehicle. **A**. Quantitation of the number of CD8^+^ T cells, CD4^+^ Tconv, and Treg in the TIL, per mg of tumor, at Day 9. **B**. Quantitation of proliferating (Ki67^+^) CD8^+^ T cells, CD4^+^ Tconv, and Treg, as a percentage of the corresponding parent population, in the TIL at Day 9. **A-B**. Data from two independent experiments (one for ATRi QDx9 group) with 2-5 mice per group. n = 7 Vehicle, 4 ATRi QDx9, 9 RT, 7 ATRi QDx9 + RT. **C**. Quantitation of proliferating (Ki67^+^) CD8^+^ T cells, CD4^+^ Tconv, and Treg, as a percentage of the corresponding parent population, in the DLN at Day 9. **D**. Quantitation of CD8^+^ TEM, as a percentage of total CD8^+^ T cells, in the Spleen and DLN at Day 9. **C-D**. Data from two independent experiments with 2-5 mice per group. n = 7 Vehicle, 7 ATRi QDx9, 8 RT (6 DLN), 7 ATRi QDx9 + RT. **E**. Representative cytograms depicting AH1 Pentamer^+^ CD8^+^ T cells in the DLN at day 9. Naïve (no tumor) negative control also shown. **F**. Quantitation of tumor antigen specific (AH1 Pentamer^+^) CD8^+^ T cells, as a percentage of total CD8^+^CD4^-^ cells, in the DLN at Day 9. Data from 5 independent experiments with 1-4 mice per group. n = 10 Vehicle, 9 ATRi QDx9, 9 RT, 12 ATRi QDx9 + RT. **A-D,F**. Mean and SD bars shown. *p<0.05, **p<0.01, ***p<0.001, ****p<0.0001 by ANOVA with Tukey’s multiple comparisons test.

Next we examined the impact of prolonged daily ATRi (QDx9) on the peripheral adaptive T cell response. We treated mice with ATRi QDx9, with or without RT, and harvested DLN and spleens on day 9. In the DLN, ATRi QDx9, independent of RT, significantly reduced the relative numbers of Treg at day 9, but did not alter the numbers of CD8^+^ T cells or CD4^+^ Tconv (Fig S7A). In spleens, ATRi QDx9 and ATRi QDx9 plus RT significantly increased the relative numbers of CD8^+^ T cells and ATRi QDx9 plus RT significantly increased the relative numbers of CD4^+^ Tconv (Fig S7B). The differential net conseqeunces of ATRi QDx9 treatment with respect to the relative sizes of T cell populations in the DLN and spleens suggest that ATRi may differentially impact other immune poulations present in these tissues. That ATRi QDx9 and ATRi QDx9 plus RT significantly decreased overall spleen weights further supports that ATRi impacts the non-T cell populations in lymphoid tissues as well (Fig S7C).

We also examined the effects of prolonged daily ATRi treatment on proliferating T cell populations in the periphery and found that ATRi QDx9, with or without RT, significantly reduced the percentages of proliferating T cells in the DLN and spleens (Fig 7C and S7D). Given this persistent suppression of proliferating T cell populations, we anticipated that ATRi QDx9 would prevent the tumor CD8^+^ TEM and antigen-specific CD8^+^ T cell expansions in the DLN that we observed after ATRi QDx3 plus RT. In contrast to short-course ATRi treatment, ATRi QDx9 and ATRi QDx9 plus RT significantly reduced the CD8^+^ TEM pools in spleens and DLN (Fig 7D), and ATRi QDx9 plus RT significantly reduced the CD8^+^ TCM pools in spleens (Fig S8A). Accordingly, the relative CD8^+^ TN pools significantly increased in spleens after ATRi QDx9, alone or with RT (Fig S8B).

Finally, we examined the DLN for tumor antigen-specific CD8^+^ T cells at day 9 (Fig 7E). We observed no significant differences across treatments in the amount of total CD8^+^CD4^-^ cells (Fig S8C) or in the percentages of pentamer^+^ CD8^+^ T cells in the DLN (Fig 7F). Therefore, prolonged daily ATRi treatment for 9 consecutive days reduces activated CD8^+^ T cell populations (TEM and TCM) in the periphery and abolishes the expansion of both the CD8^+^ TEM pool and tumor antigen-specific CD8^+^ T cells in the DLN that otherwise occurs after short-course ATRi plus RT.

## Discussion

The tumor microenvironment may be both inflammatory, as a consequence of innate immune signals and antigenic stimulation, and tolerogenic, as a consequence of the recruitment of immunosuppressive cell types including regulatory T cells and myeloid derived suppressor cells, and chronic antigen stimulation and metabolic deprivation that can cause T cell dysfunction (*60*–*66*). An established tumor contains multiple sub-populations of T cells in various degrees of function or dysfunction, including effector T cells, exhausted T cells, and regulatory T cells, that are being continuously stimulated to proliferate. The proliferation of these multiple sub-populations of T cells may be an unappreciated target of ATRi. Consistent with this premise, we show that ATRi reduces the exisiting proliferating T cell pool in both the tumor and periphery and that, with the appropriate sequence and combination treatment, this generates a niche in which new, tumor-reactive T cells can expand.

An impactful finding of our experiments is that cessation of ATRi is critical for the expansion of tumor antigen-specific CD8^+^ T cells in the periphery following treatment with short-course ATRi integrated with RT. Our data in mice show that short-course, 3-day ATRi treatment is sufficient to reduce immune populations in peripheral blood and proliferating T cell populations in TIL and lymphoid tissues. Our data are consistent with phase I data that show that ATRi suppresses circulating monocytes and proliferating T cells in patients (*31*, *32*). Cessation of this short-course, 3-day ATRi treatment results in a rapid, but nonselective, proliferative rebound in T cell populations within 4 days. When short-course ATRi treatment is integrated with RT, treatment elicits preferential expansion of tumor antigen-specific CD8^+^ T cells in the DLN. Moreover, our finding that prolonged daily ATRi treatment entirely abolishes this preferential expansion of tumor-specific CD8^+^ T cells in the DLN is consistent with the need to cease ATRi treatment to prevent restraint of the adaptive immune response. This premise is important as there are active clinical trials of immunotherapy where patients are given ATRi twice-daily for up to two weeks (*33*–*41*). Our data strongly suggest that ATRi would need to be ceased in order to generate a CD8^+^ T cell dependent response to immune checkpoint blockade. Furthermore, we find that a significant portion of the expanding tumor antigen-specific CD8^+^ T cells in the DLN have an effector/effector memory (TEM) phenotype which is consisent with our previous reports that ATRi plus RT promotes CD8^+^ T cell-dependent anti-tumor responses and immunologic memory (*17*) and may be of particular significance since recent data suggest that it is effector memory antigen-specific CD8^+^ T cells that infiltrate the tumors and drive response to immune checkpoint blockade (*67*).

Our data support that the expansion of tumor antigen-specific CD8^+^ T cells following short-course ATRi integrated with RT is driven by proinflammatory cytokine and chemokine signaling in the tumor microenvironment and downstream communication of that signaling (via immune cell migration) to the DLN. We show that both short-course and prolonged daily treatment with ATRi potentiate RT-induced IFN-β signaling in the tumor micrenvironment. Impactfully, we also show that, while short-course ATRi potentiates RT-induced increases of the IFN-β-inducible, proinflammatory chemokines CXCL10 and CCL2, prolonged ATRi treatment does not. Therefore, the duration of ATRi treatment that is integrated with RT affects the post-RT chemokine milleu in the tumor microenvironment. The dependence of both innate and adaptive immune responses after radiotherapy on IFN-β signaling in the tumor microenvironment is well established (*68*, *69*), and increases of IFN-β, CXCL10 and CCL2 in tumors affect immune cell recruitment and activation. In the short term, IFN-β signaling positively regulates NK cell function, DC maturation, and T cell activation and modulates differentiation of Ly-6C^+^ inflammatory monocytes (*70*, *71*). IFN-β also induces production of CXCL10, which recruits anti-tumor NK cells and effector T cells, and CCL2, which triggers the initial recruitment of inflammatory monocytes (*70*–*73*). Prior reports have shown that IFN-β, CXCL10, and CCL2 are increased after ATRi plus RT in the tumor microenvironment (*19*) and tumor cells cultured *in vitro* (*26*) and this has been attributed to tumor cell-intrinsic signaling. Our findings that both short-course and prolonged daily ATRi treatment potentiate RT-induced IFN-β are consistent with tumor-cell intrinsic IFN-β signaling. However, our findings that short-course but not prolonged daily ATRi treatment potentiate RT-induced CXCL10 and CCL2 suggest another source for these chemokines. In response to proinflammatory stimuli, CXCL10 can be secreted by T cells and monocytes and CCL2 can be secreted by myeloid lineage cells including monocytes, macrophages, and DC (*70*, *72*, *74*). Therefore, it is possible that tumor-infiltrating myeloid cells are an important source of CXCL10 and CCL2 after short-course ATRi treatment integrated with RT, and that the negative impact of prolonged ATRi treatment on these myeloid cell populations suppresses the production of these proinflammatory chemokines.

Consistent with increased proinflammatory signaling in the tumor microenvironment, our data suggest that short-course, 3-day ATRi treatment integrated with RT increases migration of inflammation-associated cells, including NK cells, CD11b^+^CD11c^+^ cells, and CD11c^+^CD11b^+^Ly-6C^+^ cells, to the tumor-draining lymph node (DLN). CD11b^+^CD11c^+^ cells include both CD11b^+^ dendritic cells and CD11c-expressing monocytes. Ly-6C-expressing CD11c^+^CD11b^+^ cells arise at inflamed tissues and reside along a differentiation spectrum between inflammatory monocytes and antigen-presenting, monocyte-derived dendritic cells (moDC) (*48*, *51*–*57*). Inflammatory monocytes can phagocytose antigens, migrate to the lymph node, and differentiate into antigen-presenting moDC, which then present antigen to CD4^+^ T cells, cross-present antigen to CD8^+^ T cells, or transfer antigen to lymph node-resident DC, which, in turn, cross-present to CD8^+^ T cells (*48*, *51*, *53*, *55*, *75*–*77*). Preclinincal evidence supports an essential role for moDC that accumulate in the tumor and DLN in the activation of CD8^+^ T cells and anti-tumor immunity after immunotherapy (*57*). Our data do not definitively demonstrate that the CD11c^+^CD11b^+^Ly-6C^+^ pool in the DLN represents differentiated, antigen-presenting moDC that are responsible for the increased priming of CD8^+^ T cells in the DLN after short-course, 3-day ATRi treatment integrated with RT. However, our data do associate accumulation of these cells in the inflamed DLN with increased CD8^+^ T cell activation at day 7 and subsequent tumor antigen-specific CD8^+^ T cell expansion at day 9.

To summarize, our work herein highlights the importance of proper scheduling of ATRi for immune modulation. We associate the expansion of a tumor antigen-specific CD8^+^ T cells after short-course, 3-day ATRi treatment integrated with RT with both the potentiation of RT-induced inflammation and innate immune activity by ATRi as well as the independent effects of ATRi on T cells and the proliferative rebound that follows cessation of ATRi. Despite potentiation of RT-induced IFN-β in the tumor microenvironment, prolonged daily ATRi treatment restrains the peripheral adaptive T cell response leaving the tumor infiltrate almost devoid of T cells and entirely abolishes the peripheral expansion of tumor antigen-specific effector CD8^+^ T cells that otherwise occurs after short-course ATRi treatment integrated with RT. Therefore, we demonstrate that cessation of ATRi is essential to allow for peripheral T cell rebound and the expansion of tumor-specific CD8^+^ T cell clones. Our data suggest cessation of ATRi may be important for the innate immune response as well. Whereas short-course ATRi potentiated RT-induced production of some proinflammatory chemokines in the tumor microenvironment, prolonged ATRi treatment negated this potentiation. These findings have important implications for the clinical deployment of ATRi as part of regimens where expansion of tumor antigen-specific effector CD8^+^ T cells is desirable, such as immunotherapy, as improper schedule of ATRi treatment may restrain the adaptive immune system and negatively impact the efficacy of immunotherapy. Our findings also have relevance to the numerous combinations of ATRi with cytotoxic backbones currently under clinical investigation, which have been shown to induce immunogenic cell death (*1*, *78*, *79*). Prolonged exposure to ATRi may well limit the immunogenic response and thereby the therapeutic benefit that otherwise might be achieved in patients.

## Materials and Methods

### Cell lines and reagents

CT26 (ATCC CRL-2638) were purchased from ATCC and cultured in RPMI containing 10 % FBS, 100 U/mL penicillin, and 100 mg/mL streptomycin (all Lonza). Cells were routinely tested for mycoplasma. AZD6738 (ATRi) was provided by AstraZeneca and dosed at 75 mg/kg by oral gavage as previously described (*15*, *17*).

### Mice and treatments

Female BALB/C (6-8 weeks old) were purchased from Jackson Laboratories. CT26 cells (~5 × 10^5^) in RPMI were subcutaneously injected into the right hind flank of approximately 8–10-week-old mice. Treatment was initiated 7-10 days post injection when tumors reached ~60-120 mm^3^. For tumor irradiation, immobilized mice received two fractions of 2 Gy (6 mV photon energy, 2 cm field) on days 1-2. Mice received 75 mg/kg AZD6738 approximately 45 min prior to irradiation on days 1-2 and a third dose, approximately 18 hours later, on day 3. Subsequent doses of AZD6738 for prolonged daily treatment experiments were given daily thereafter, approximately every 24 hours.

### Tissue processing, staining, and multi-parameter flow cytometry for immunoprofiling

CT26 tumors, tumor-draining (right inguinal) lymph nodes (DLN), and spleens were harvested from mice at days 4, 7, and 9. Tissues were processed to generate single cell suspensions, as previously described (*17*). Briefly, DLN and spleens were mechanically dissociated between frosted glass slides and filtered through 70 μm cell strainers (Corning). Tumors were injected in multiple sites with a total volume of 1.5 mL Liberase enzyme digestion solution (50 μg/mL Liberase DL research grade (Roche) and 10 U/mL DNase I (Sigma) in RPMI) and incubated 3 min at room temperature. Tumors were then cut into small pieces and incubated in a total volume of 5 mL Liberase DL/DNase solution for 15 min at 37°C. After enzymatic digestion, tumors were mechanically dissociated between frosted glass slides, filtered through 70 μm cell strainers (Corning), vortexed at low speed for 90 sec, and filtered again through new 70 μm cell strainers (Corning). Erythrocytes were lysed in 150 mM NH4Cl, 10 mM NaHCO3, 0.1 mM EDTA pH 8.0. for 10 sec (tumors) or 30 sec (spleens). Single cell suspensions were counted using a Millipore Scepter 2.0 or 3.0 handheld counter (Millipore) or Cellometer K2 (Nexcelom) and seeded at 1.5-2 × 10^6^ cells (equivalent density within a given experiment) in 96-well round bottom plates for staining. Cells were then blocked in FSC buffer (2% FBS/1x PBS) containing 0.5 μg anti-CD16/32 antibody (TruStain FcX Plus, BioLegend) for 10 min at 4°C to block non-specific binding of antibodies via Fc receptors. For tumor antigen specific CD8^+^ T cell experiments, cells were stained in FSC buffer containing 10 μL PE-labeled H-2Ld SPSYVYHQF (AH1 peptide) Pro5 MHC Class I Pentamer (ProImmune) for 20 min at room temperature. Lymph node from a naïve (no tumor) negative control mouse was included with each experiment, and for AZD6738 QDx9 experiments, PE-labeled HLA-A*02:01 Negative Control Pro5 MHC Class I Pentamer (ProImmune) was also included. Cells were then stained in FSC buffer containing antibodies to surface antigens for 15 min at 4°C. For staining panels including multiple Brilliant Violet dye-conjugated antibodies, Brilliant Stain Buffer Plus (BD Biosciences) was included in the staining cocktail to prevent polymer dye-dye interactions. For staining panels including PE-tandem and PerCP-tandem dye-conjugated antibodies, True-Stain Monocyte Blocker (BioLegend) was included in the staining cocktail to prevent non-specific binding of monocytes and macrophages to the tandem dyes. Both Brilliant Stain Buffer Plus and True-Stain Monocyte Blocker were used according to the manufacturer’s instructions. Following surface staining, cells were stained with eFlour780 viability dye (1:4000, ThermoFisher) in 1x PBS for 10 minutes at 4°C to label dead/dying cells, fixed and permeabilized in eBioscience Fixation/Permeabilization reagent (ThermoFisher) for 15 min at room temperature, and when performing nuclear staining (Ki67, Foxp3), stained for 45 min at room temperature in eBioscience 1x Permeabilization Buffer (ThermoFisher) containing antibodies to nuclear proteins. Uncompensated data were collected using a BD LSRFortessa 4-laser cytometer and BD FACSDiva software. Compensation and data analyses were performed in FlowJo V10 software. Single stained spleen or DLN samples with matching unstained cells or single stained OneComp eBeads (ThermoFisher) were used for single color compensation controls. Fluorescence-minus-one (FMO) controls were used, where appropriate, to empirically determine gating. Antibodies for staining panels are included in Table S1. Gating strategies are shown in Figures S9-S13.

### Measurement of intratumoral cytokines and chemokines

Tumors were harvested at the indicated timepoints, cut into smaller pieces, and snap-frozen on dry ice. Frozen tumor pieces were weighed and approximately 50-60 mg of tumor tissue was added to a Precellys CK28-R Protein Safe Hard tissue homogenizing tube (Bertin Technologies) containing Invitrogen ProcartaPlex Cell Lysis Buffer (ThermoFisher) with 1mM PMSF, at a volume of 500 μL lysis buffer per 75 mg tumor tissue. Homogenization and protein extraction were performed using a Precellys 24 homogenizer (Bertin Instruments) with the following protocol: 2 cycles of 6000 RPM x 15 sec, samples on ice ≥2 min, 2 additional cycles of 6000 RPM x 15 sec. Tumor extracts were transferred to new tubes and cleared of insoluble material prior to freezing aliquots at −80°C. Amounts of the following 10 protein targets in the tumor extracts were measured using a customized 10-plex U-Plex Assay (Mesoscale Discovery): GM-CSF, IFN-α, IFN-β, IFN-γ, IL-6, IL-12p70, CXCL10 (IP-10), CCL2 (MCP-1), CCL3 (MIP-1α), and CCL5 (RANTES). The assay was performed according to the manufacturer’s instructions, with the modification that, after coating the plate, samples were incubated at room temperature with shaking for 1 hour and then at 4°C overnight prior to proceeding to the antibody detection step. For GM-CSF, IFN-α, IFN-β, IFN-γ, IL-6, IL-12p70, and CCL5 (RANTES), the assay was performed using undiluted tumor extract. For CXCL10 (IP-10), CCL2 (MCP-1), and CCL3 (MIP-1α), tumor extracts were diluted 1:50 in the diluent provided by the manufacturer. All samples were assayed in duplicate. Assay detection was performed using a MESO QuickPlex SQ 120 instrument (Mesoscale Discovery). Raw data analyses and determination of protein concentrations (in pg/mL) in the tumor extracts were performed using MSD Discovery Workbench software (Mesoscale Discovery). Protein concentrations were normalized to the mean concentrations of vehicle control samples for a given target, assayed on the same plate, and the data are presented as the relative amount of protein compared to vehicle control. No inter-plate or inter-run comparisons were made.

### Statistics

For CBC experiments, statistical significance was determined by two-tailed, unpaired t test (95% confidence interval). For all other experiments, statistical significance was determined by ANOVA with Tukey’s multiple comparison tests (95% confidence level). Data are reported as mean ± SD. Brackets are shown only for comparisons that were statistically significant. All statistical analyses were performed in Graphpad Prism 9.

### Animal Study approval

Experimental procedures were approved by the University of Pittsburgh Animal Care and Use Committees and performed in accordance with relevant guidelines and regulations.

## Supporting information

Vendetti Supplemental

## Acknowledgements

We thank Mark J. O’Connor, PhD (AstraZeneca) for providing AZD6738 and Ron LaLonde, PhD, DABR (UPMC Radiation Physicist) for technical assistance with our experiments. This work was supported by R01CA236367 and R01CA204173 (CJB). This project used the Animal Facility, Cancer Pharmacokinetics and Pharmacodynamics Facility, and the Cytometry Facility that are supported in part by award P30CA047904 from the NIH.

## Author contributions

FPV, DAC, JHB, and CJB designed experiments. FPV, SS, MD, NI, and JC executed experiments. FPV and CJB analyzed experiments. FPV, DAC, JHB, and CJB wrote the manuscript.

