## Supplementary material for "Short-course but not prolonged treatment with ATR inhibitor AZD6738 integrates with radiotherapy to generate a tumor antigen-specific CD8^+^ T cell expansion in the periphery": Vendetti Supplemental

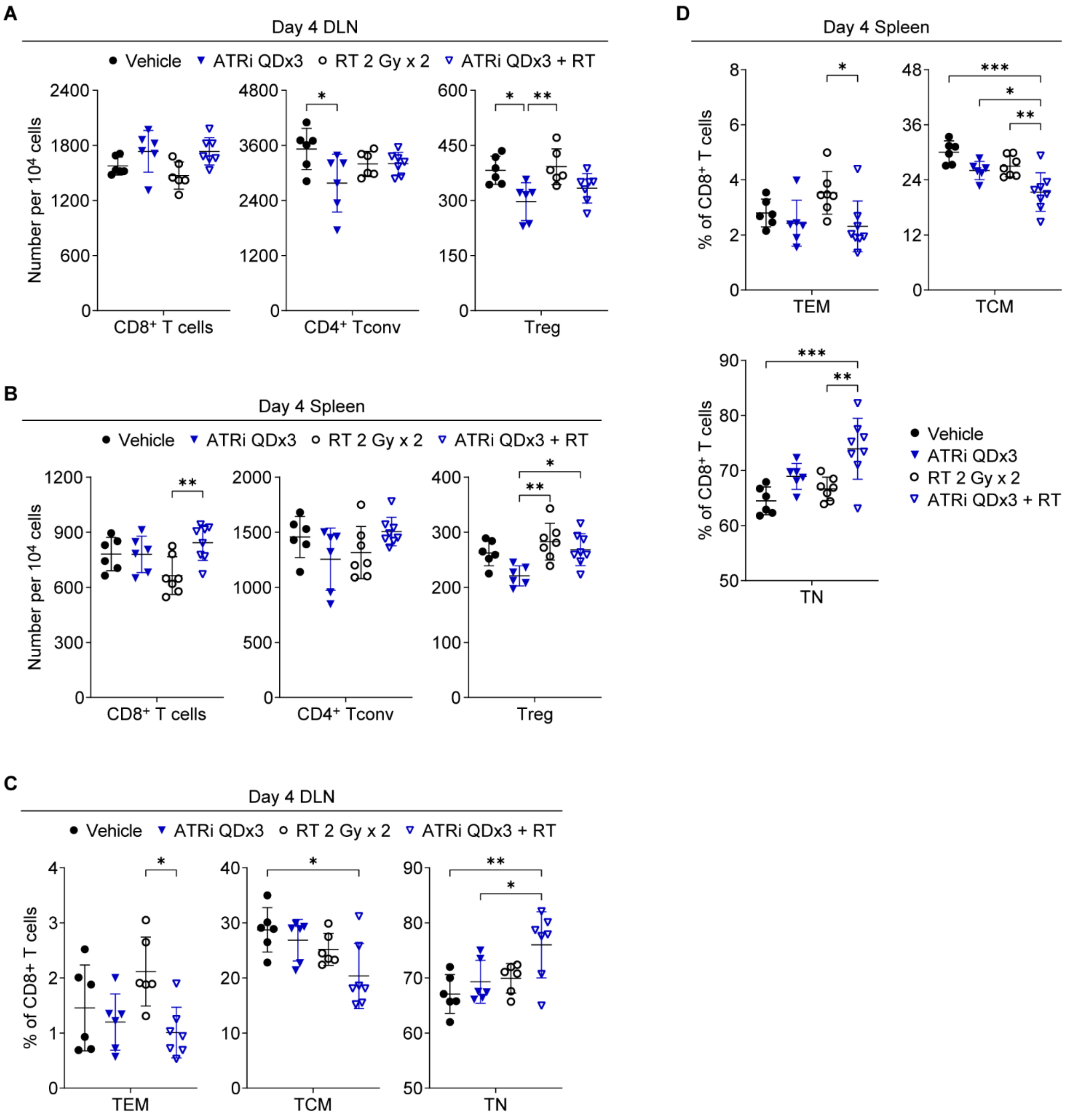


**Figure S1. Effects of short-course ATRi treatment in the TIL and periphery at Day 4.**

**A-D.** CT26 tumor-bearing mice were treated with ATRi on days 1-3 (ATRi QDx3), radiotherapy on days 1-2 (RT 2 Gy x 2), ATRi QDx3 + RT, or vehicle. **A.** Quantitation of the relative number of $CD8^+$ T cells, $CD4^+$ Tconv, and Treg, per $10^4$ cells stained, in the DLN at Day 4. **B.** Quantitation of the relative number of $CD8^+$ T cells, $CD4^+$ Tconv, and Treg per $10^4$ cells stained, in spleens at Day 4. **C.** Quantitation of $CD8^+$

TEM, TCM, and TN, as a percentage of total $CD8^+$ T cells, in the DLN at day 4. **D.** Quantitation of $CD8^+$ TEM, TCM, and TN, as a percentage of total $CD8^+$ T cells, in spleens at Day 4. **A-D**. Data from 3 independent experiments with 1-3 mice per group. n = 6 Vehicle, 6 ATRi QDx3, 7 RT (6 DLN), 8 ATRi QDx3 RT (7 DLN). Mean and SD bars shown. *p<0.05, **p<0.01, ***p<0.001 by ANOVA with Tukey's multiple comparisons test.

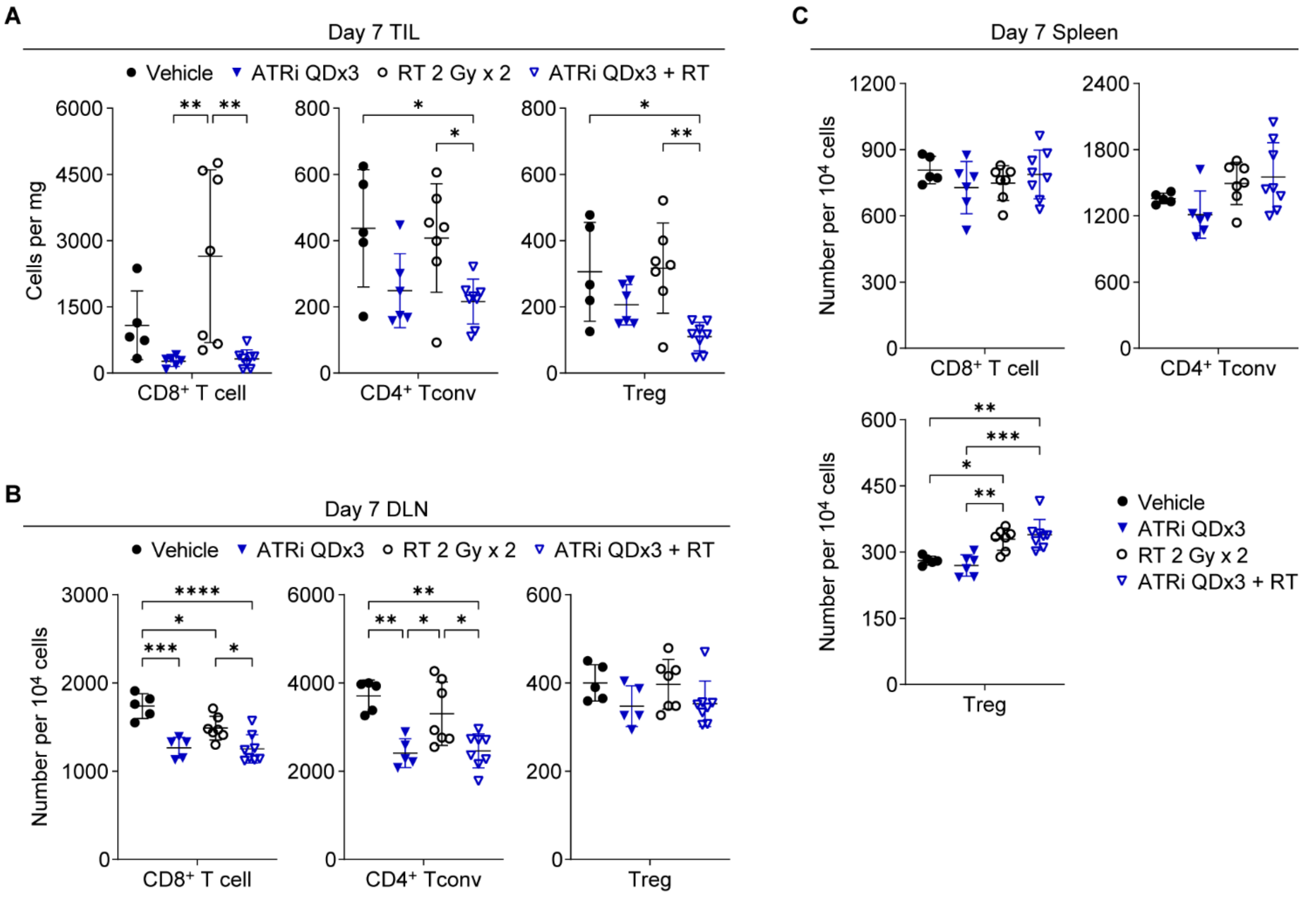


**Figure S2. Cessation of short-course ATRi treatment results in a proliferative rebound in T cells by Day 7.**

**A-E.** CT26 tumor-bearing mice were treated with ATRi QDx3, RT (2 Gy x 2), ATRi QDx3 + RT, or vehicle. **A.** Quantitation of the number of $CD8^+$ T cells, $CD4^+$ Tconv, and Treg, per mg of tumor stained, in the TIL at Day 7. **B.** Quantitation of the relative number of $CD8^+$ T cells, $CD4^+$ Tconv, and Treg, per $10^4$ cells stained, in the DLN at Day 7. **C.** Quantitation of the relative number of $CD8^+$ T cells, $CD4^+$ Tconv, and Treg, per $10^4$ cells stained, in spleens at Day 7. **A-C**. Data from at least 2 independent experiments with 1-4 mice per group. n = 5 Vehicle, 6 ATRi QDx3 (5 DLN), 7 RT, 8 ATRi QDx3 + RT. Mean and SD bars shown. *$p<0.05$, **$p<0.01$, ***$p<0.001$, ****$p<0.0001$ by ANOVA with Tukey's multiple comparisons test.

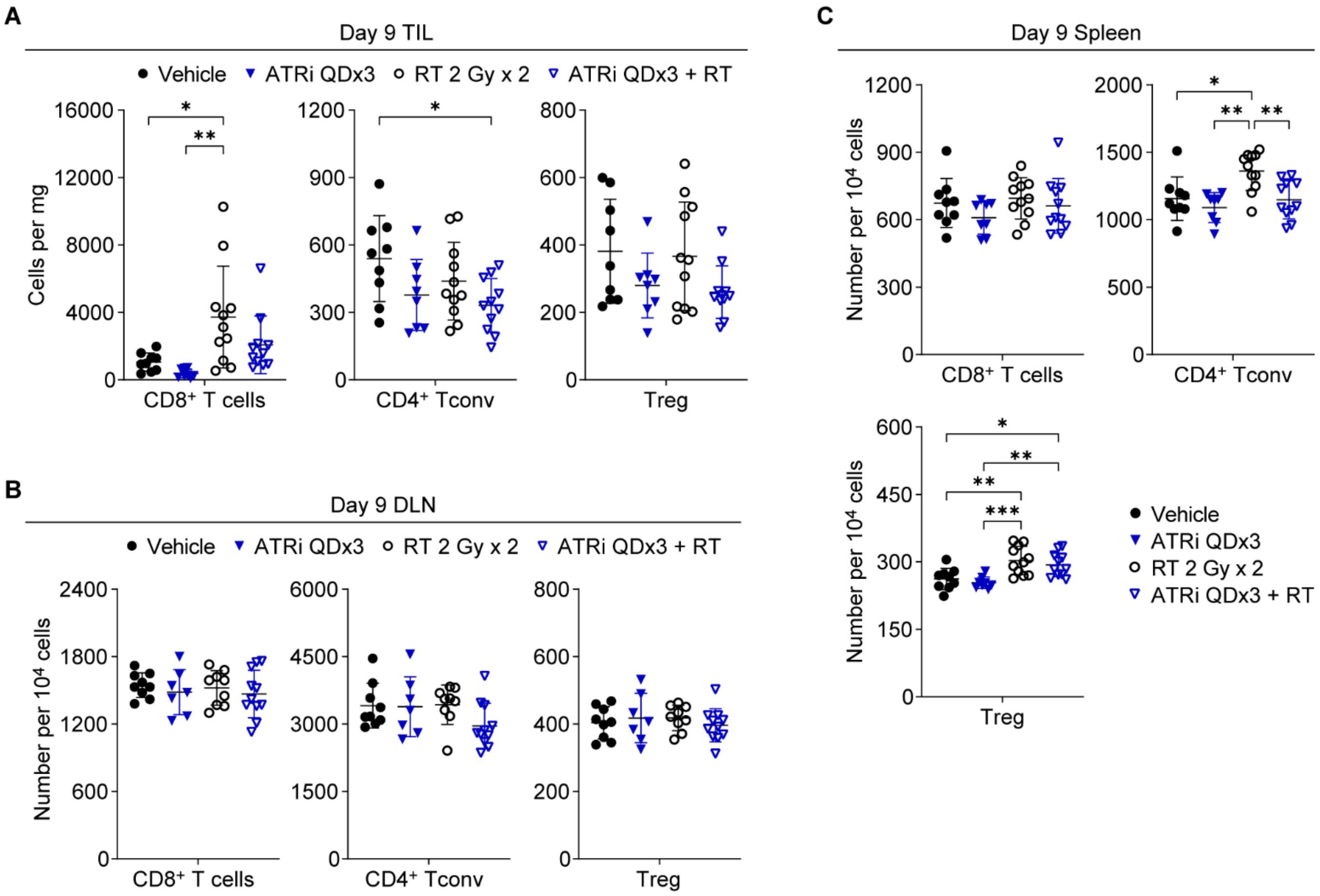


**Figure S3. Repopulation of T cells by Day 9 in the TIL and periphery after cessation of short-course ATRi treatment.**

**A-C.** CT26 tumor-bearing mice were treated with ATRi QDx3, RT (2 Gy x 2), ATRi QDx3 + RT, or vehicle. **A.** Quantitation of the number of $CD8^+$ T cells, $CD4^+$ Tconv, and Treg, per mg of tumor stained, in the TIL at Day 9. **B.** Quantitation of the relative number of $CD8^+$ T cells, $CD4^+$ Tconv, and Treg, per $10^4$ cells stained, in the DLN at Day 9. **C.** Quantitation of the relative number of $CD8^+$ T cells, $CD4^+$ Tconv, and Treg, per $10^4$ cells stained, in spleens at Day 9. **A-C.** Data from at least 4 independent experiments with 1-3 mice per group. n = 9 Vehicle, 8 ATRi QDx3 (7 DLN), 11 RT (9 DLN), 11 ATRi QDx3 + RT. Mean and SD bars shown. *p<0.05, **p<0.01, ***p<0.001 by ANOVA with Tukey's multiple comparisons test.

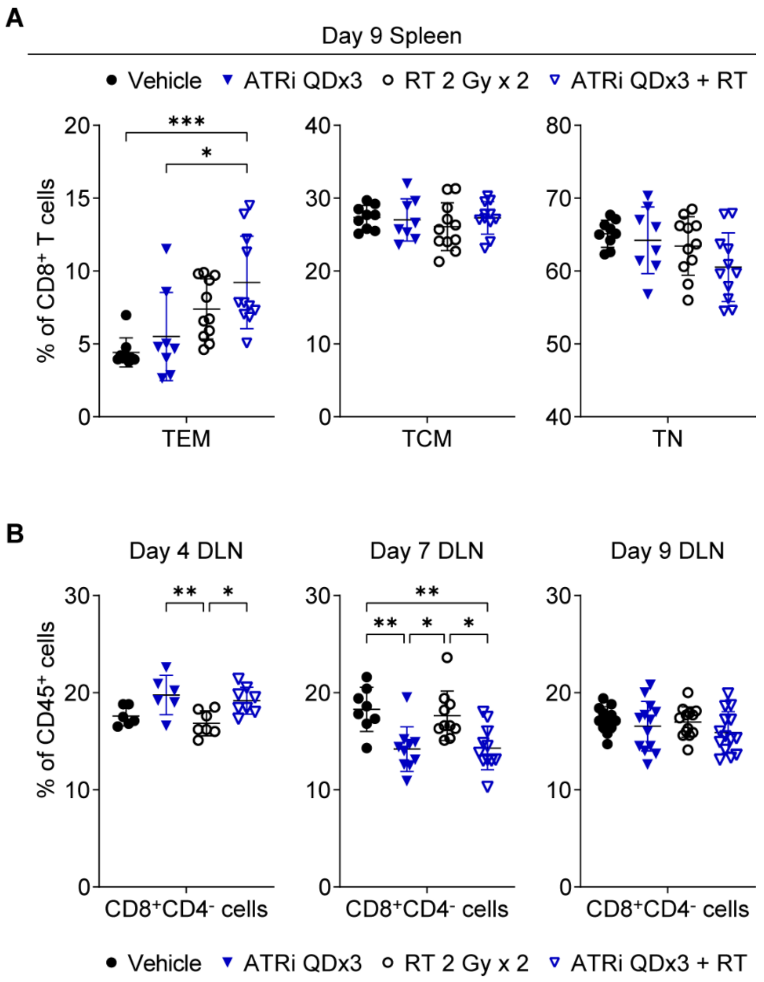


**Figure S4. Short-course ATRi integrated with RT promotes expansion of effector T cells in the periphery.**

**A-B.** CT26 tumor-bearing mice were treated with ATRi QDx3, RT (2 Gy x 2), ATRi QDx3 + RT, or vehicle. **A.** Quantitation of $CD8^+$ TEM, TCM, and TN, as a percentage of total $CD8^+$ T cells, in spleens at day 9. Data from at least 4 independent experiments with 1-3 mice per group. n = 9 Vehicle, 8 ATRi QDx3, 11 RT, 11 ATRi QDx3 + RT. **B.** Quantitation of $CD8^+CD4^-$ cells, as a percentage of total $CD45^+$ cells, in the DLN at Day 4, Day 7, and Day 9. Data from 3 (Day 4), at least 3 (Day 7), and at least 5 (Day 9) independent experiments with 1-5 mice per group. n at Day 4 = 6 Vehicle, 6 ATRi QDx3, 7 RT, 8 ATRi QDx3 + RT. n at Day 7 = 8 Vehicle, 10 ATRi QDx3, 10 RT, 11 ATRi QDx3 + RT. n at Day 9 = 12 Vehicle, 13 ATRi QDx3, 13 RT, 14 ATRi QDx3 + RT. **A-B**. Mean and SD bars shown. *p<0.05, **p<0.01, ***p<0.001, ****p<0.0001 by ANOVA with Tukey's multiple comparisons test.

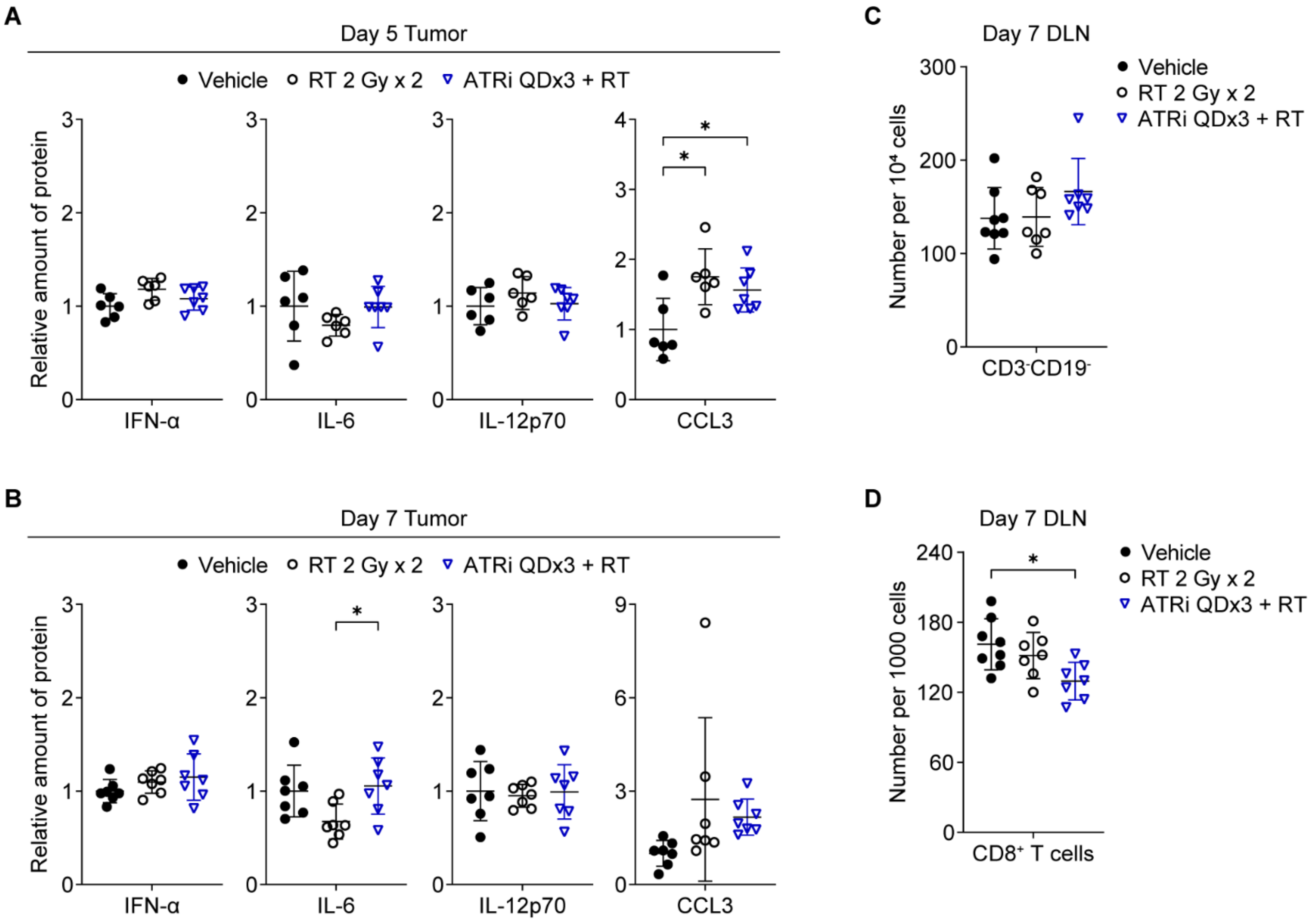


**Figure S5. Impact of short-course ATRi treatment on proinflammatory cytokines and chemokines in the tumor microenvironment.**

**A-D.** CT26 tumor-bearing mice were treated with RT (2 Gy x 2), ATRi QDx3 + RT, or vehicle. **A.** Quantitation of the relative amount of protein (normalized to vehicle control) of a subset of inflammatory cytokines and chemokines in tumors at Day 5. **B.** Quantitation of the relative amount of protein (normalized to vehicle control) of a subset of inflammatory cytokines and chemokines in tumors at Day 7. **A,B.** Data from one experiment (Day 5) or two independent experiments (Day 7). n at Day 5 = 6 Vehicle, 6 RT, 7 ATRi QDx3 + RT. n at Day 7 = 7 Vehicle, 7 RT, 7 ATRi QDx3 + RT. **C.** Quantitation of the relative number of $CD3^-CD19^-$ cells in the DLN at Day 7. **D.** Quantitation of the relative number of $CD8^+$ T cells in the DLN at Day 7. **C,D.** Data from two independent experiments with 3-4 mice per group. n = 8 Vehicle, 7 RT, 7 ATRi QDx3 + RT. **A-D**. Mean and SD bars shown. *p<0.05 by ANOVA with Tukey's multiple comparisons test.

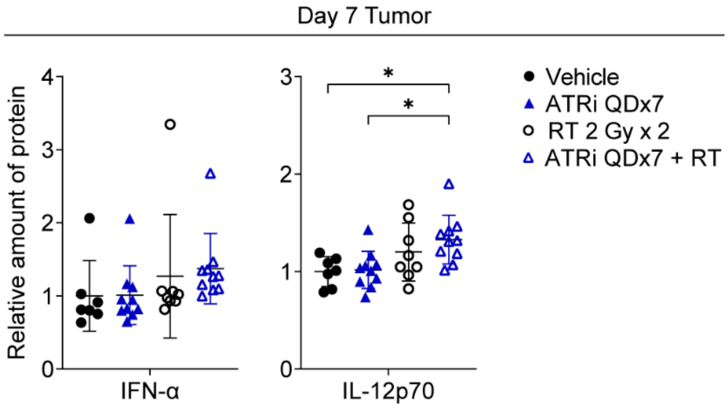


**Figure S6. Impact of prolonged daily ATRi treatment on proinflammatory cytokines in the tumor microenvironment.**

CT26 tumor-bearing mice were treated with ATRi on days 1-7 (ATRi QDx7), radiotherapy on days 1-2 (RT 2 Gy x 2), ATRi QDx7 + RT, or vehicle. Shown is the quantitation of the relative expression (compared to vehicle control) of IFN-α and IL-12p70 in tumors at Day 7. Data from at least 4 independent experiments with 1-4 mice per group. n = 7 Vehicle, 10 ATRi QDx7, 8 RT, 10 ATRi QDx7 + IR. Mean and SD bars shown. *$p<0.05$ by ANOVA with Tukey's multiple comparisons test.

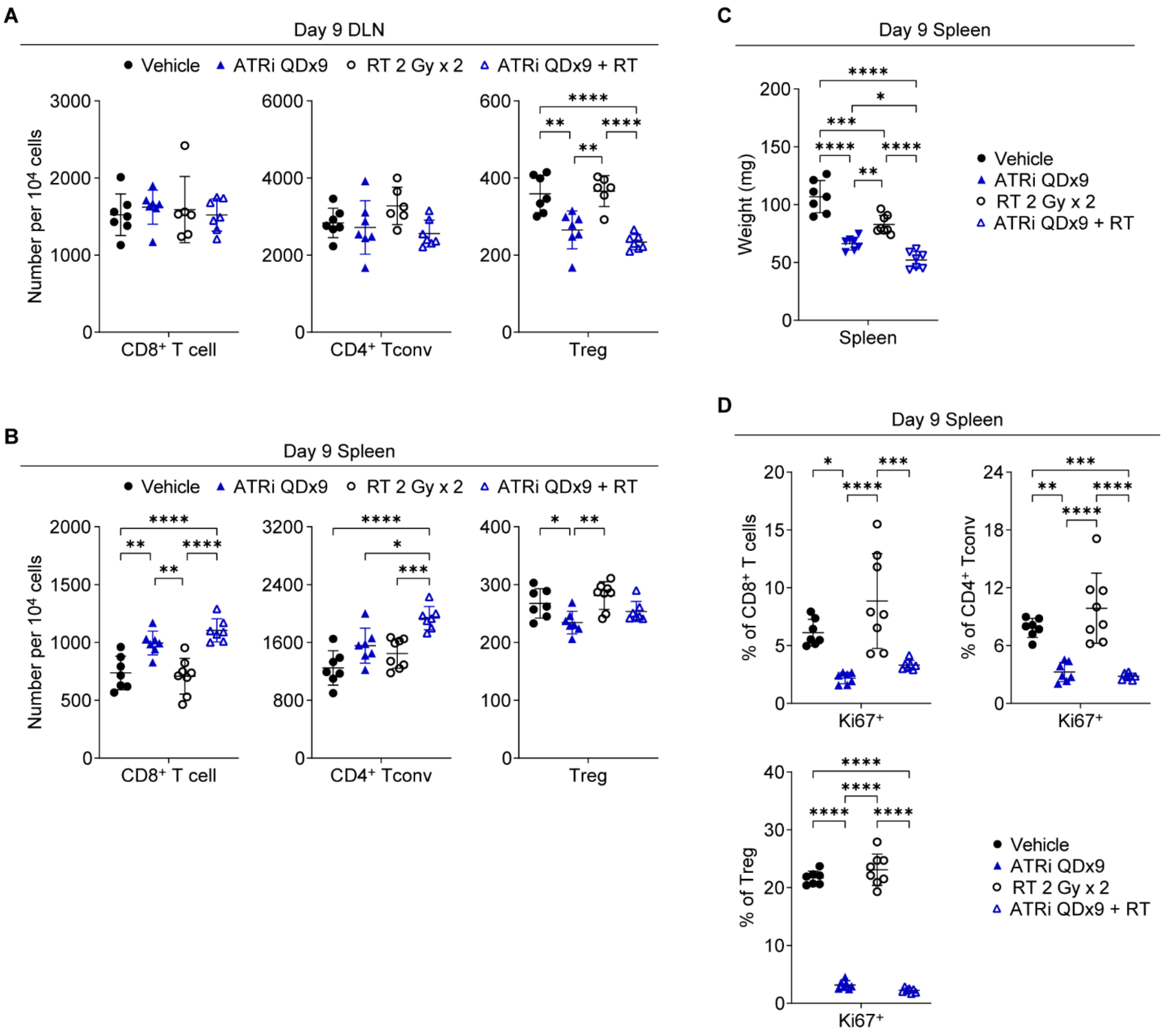


**Figure S7. Prolonged daily ATRi treatment restrains the adaptive T cell response in the periphery.**

**A-C.** CT26 tumor-bearing mice were treated with ATRi on days 1-9 (ATRi QDx9), radiotherapy on days 1-2 (RT 2 Gy x 2), ATRi QDx9 + RT, or vehicle. **A.** Quantitation of the relative number of $CD8^+$ T cells, $CD4^+$ Tconv, and Treg in the DLN at Day 9. **B.** Quantitation of the relative number of $CD8^+$ T cells, $CD4^+$ Tconv, and Treg in spleens at Day 9. **C.** Quantitation of spleens weights at Day 9. **D.** Quantitation of proliferating ($Ki67^+$) $CD8^+$ T cells, $CD4^+$ Tconv, and Treg, as a percentage of the corresponding parent population, in spleens at Day 9. **A-D.** Data from two independent experiments with 2-5 mice per group. n = 7 Vehicle, 7 ATRi QDx9, 8 RT (6 DLN), 7 ATRi QDx9 + RT. Mean and SD bars shown. *$p<0.05$, **$p<0.01$, ***$p<0.001$, ****$p<0.0001$ by ANOVA with Tukey's multiple comparisons test.

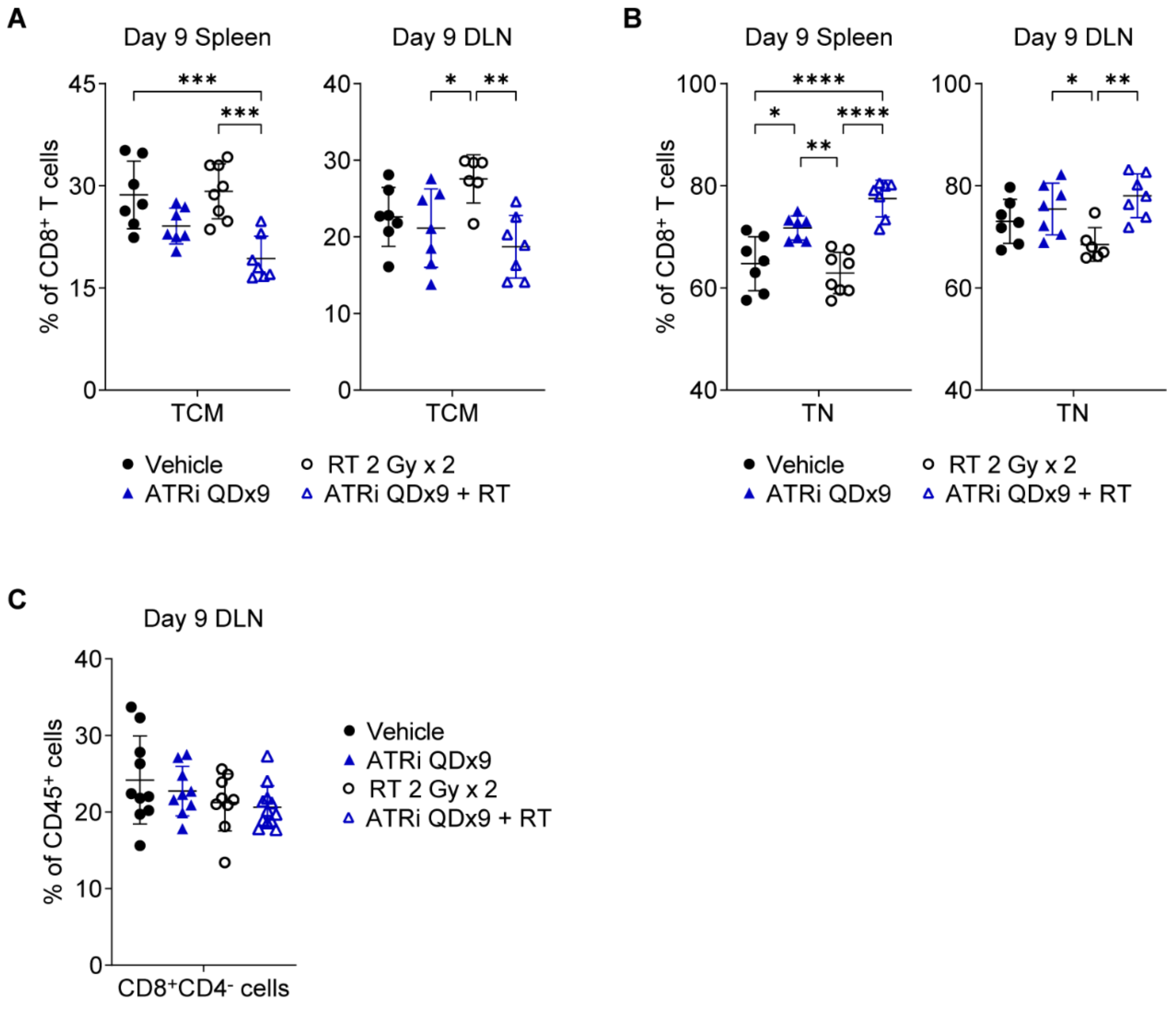


**Figure S8. Prolonged ATRi treatment restrains the adaptive CD8$^+$ T cell response in the periphery following RT.**

**A-C.** CT26 tumor-bearing mice were treated with ATRi QDx9, RT (2 Gy x 2), ATRi QDx9 + RT, or vehicle. **A.** Quantitation of CD8$^+$ TCM, as a percentage of total CD8$^+$ T cells, in the Spleen and DLN at Day 9. **B.** Quantitation of CD8$^+$ TN, as a percentage of total CD8$^+$ T cells, in the Spleen and DLN at Day 9. **A-B.** Data from two independent experiments with 2-5 mice per group. n = 7 Vehicle, 7 ATRi QDx9, 8 RT (6 DLN), 7 ATRi QDx9 + RT. **C.** Quantitation of CD8$^+$CD4$^-$ cells, as a percentage of total CD45$^+$ cells, in the DLN at Day 9. Data from 5 independent experiments with 1-4 mice per group. n = 10 Vehicle, 9 ATRi QDx9, 9 RT, 12 ATRi QDx9 + RT. **A-C.** Mean and SD bars shown. *$p<0.05$, **$p<0.01$, ***$p<0.001$ by ANOVA with Tukey's multiple comparisons test.

| Antibody | Dilution | Manufacturer | Catalog # |
|---|---|---|---|
| **Brilliant Stain Buffer Plus** | 1:5 | BD Biosciences | 566385 |
| **BV786 anti-mouse CD62L (clone MEL-14)** | 1:500, 1:1000 | BD Biosciences | 564109 |
| **PE-CF594 anti-mouse CD25 (clone PC61)** | 1:500 | BD Biosciences | 562695 |
| **AF488 anti-mouse CD3 (clone 17A2)** | 1:250 | BioLegend | 100212 |
| **AF488 anti-mouse CD8α (clone 53-6.7)** | 1:250 | BioLegend | 100723 |
| **AF488 anti-mouse CD19 (clone 6D5)** | 1:250 | BioLegend | 115524 |
| **AF488 anti-mouse CD45 (clone 30-F11)** | 1:500 | BioLegend | 103122 |
| **AF488 anti-mouse TCR-β (clone H57-597)** | 1:250 | BioLegend | 109215 |
| **AF647 Rat IgG2a,κ Isotype Control (clone RTK2758)** | 1:200 | BioLegend | 400526 |
| **AF647 anti-mouse Ki67 (clone 16A8)** | 1:200 | BioLegend | 652408 |
| **AF647 anti-mouse Ly-6C (clone HK1.4)** | 1:500 | BioLegend | 128010 |
| **BV421 anti-mouse CD69 (clone H1.2F3)** | 1:250 | BioLegend | 104545 |
| **BV421 anti-mouse NKp46 (CD335) (clone 29A1.4)** | 1:50 | BioLegend | 137611 |
| **BV510 anti-mouse CD4 (clone GK1.5)** | 1:250, 1:500 | BioLegend | 100449 |
| **BV785 anti-mouse CD11c (clone N418)** | 1:125, 1:250 | BioLegend | 117336 |
| **BV785 anti-mouse CD127 (IL-7Rα) (clone A7R34)** | 1:100 | BioLegend | 135037 |
| **PE anti-mouse/human CD11b (clone M1/70)** | 1:250 | BioLegend | 101207 |
| **PE anti-mouse/human KLRG1 (MAFA) (clone 2F1/KLRG1)** | 1:250 | BioLegend | 138407 |
| **PE-Cy7 anti-mouse CD8α (clone 53-6.7)** | 1:500 | BioLegend | 100721 |
| **PE-Cy7 anti-mouse TCRβ (clone H57-597)** | 1:250 | BioLegend | 109221 |
| **PerCP-Cy5.5 anti-mouse/human CD44 (clone IM7)** | 1:500 | BioLegend | 103031 |
| **PerCP-Cy5.5 anti-mouse CD103 (clone 2E7)** | 1:50 | BioLegend | 121416 |
| **TruStain FcX PLUS anti-mouse CD16/32 (clone S17011E)** | 1:100 | BioLegend | 156604 |
| **True-Stain Monocyte Blocker** | 1:20 | BioLegend | 426103 |
| **eFluor450 anti-mouse/rat Foxp3 (clone FJK-16s)** | 1:200 | Invitrogen | 48-5773-82 |
| **AF647 anti-mouse CD8 (clone KT15)** | 1:500 | MBL Intl | D271-A64 |
| **PE-labelled Pro5 MHC I Pentamer (H-2Ld SPSYVYHQF)** | 10 μL/well | ProImmune | F398-2A/2B |
| **PE-labelled Pro5 MHC I Pentamer (A*02:01 Negative)** | 10 μL/well | ProImmune | FN01-2A |

**Table S1. Antibodies for flow cytometry immunoprofiling**

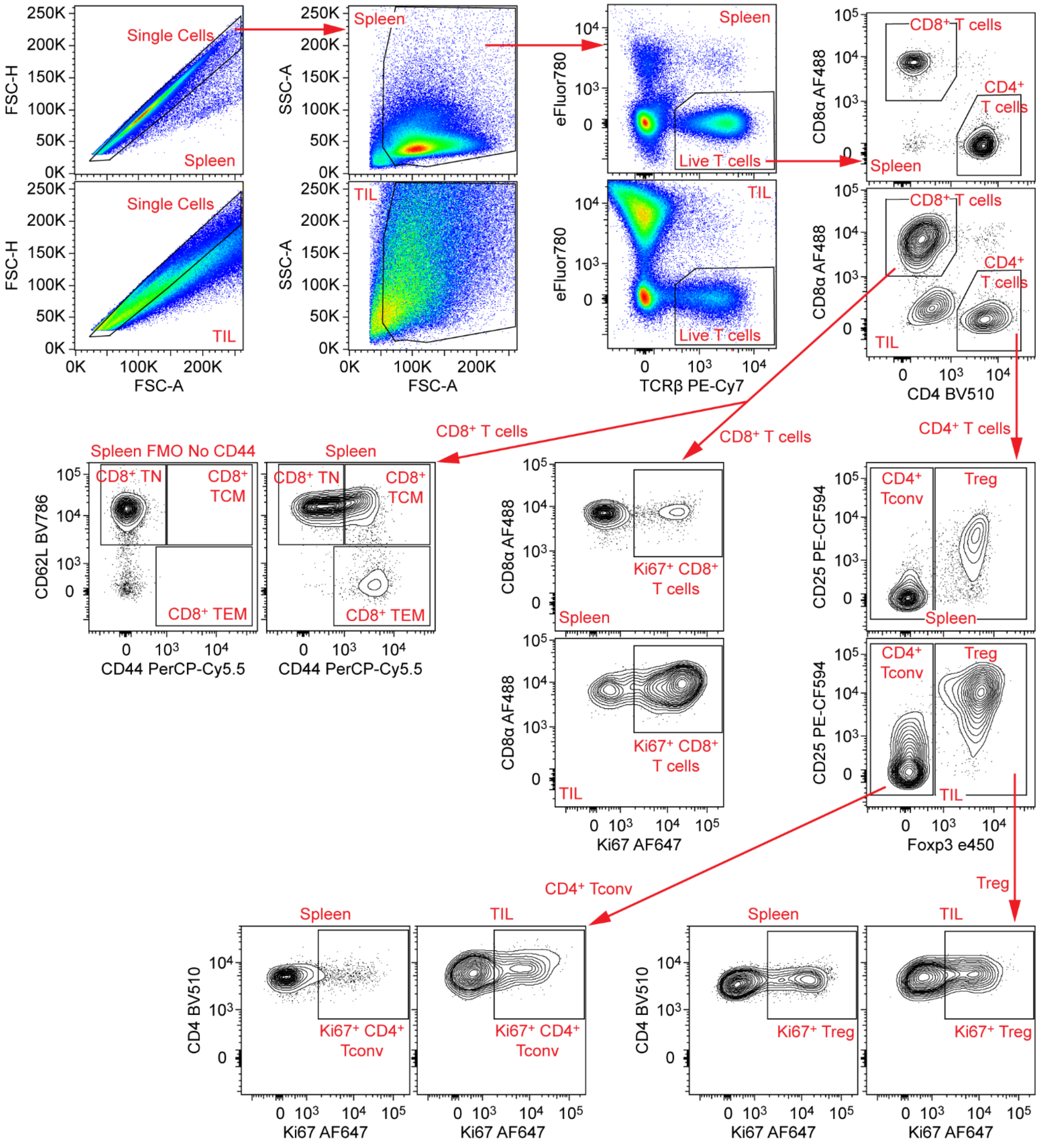


**Figure S9. Flow cytometry gating strategy to identify T cells, proliferating T cells, and CD8⁺ T cell activation phenotypes in the TIL, DLN, and spleens.**

Any unstable portions of the run were gated out prior to analysis. After single cell gating (FSC-A vs. FSC-H) to remove doublets/clumps and scatter gating (FSC-A vs. SSC-A) to exclude debris and, live T cells (efluor780$^{neg}$ and TCRβ$^{+}$) were gated and then subset into CD8$^{+}$ and CD4$^{+}$ T cells. CD8$^{+}$ T cells were examined for Ki67 expression to identify proliferating (Ki67$^{+}$) CD8$^{+}$ T cells as well as expression of CD62L and CD44 to identify naïve (CD62L$^{hi}$CD44$^{lo}$), central memory (TCM, CD62L$^{hi}$CD44$^{hi}$), and effector/effector memory (TEM, CD62L$^{lo}$CD44$^{hi}$) CD8$^{+}$ T cells. CD4$^{+}$ T cells were subset into conventional CD4$^{+}$ T cells (Tconv, Foxp3$^{neg}$) and regulatory T cells (Treg, Foxp3$^{+}$). CD25 was included to improve resolution between the CD4$^{+}$ T conv and Treg populations. Both the CD4$^{+}$ Tconv and Treg populations were then examined for Ki67 expression to identify proliferating (Ki67$^{+}$) cells in each pool. Spleen and TIL examples are shown for gating. A spleen fluorescence-minus-one control is shown for empiric determination of the CD44$^{hi}$ gate.

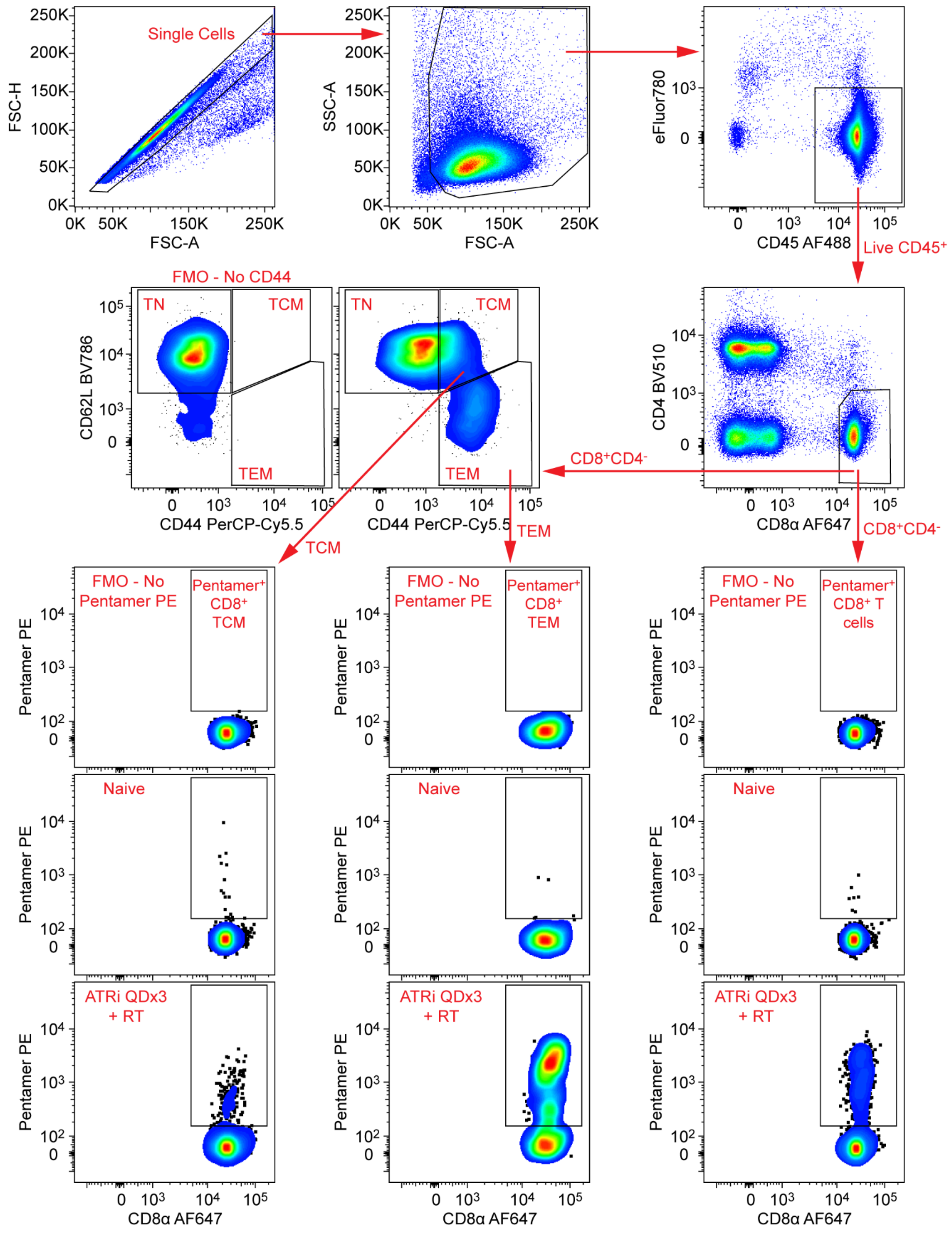

Single Cells
FSC-H
FSC-A
SSC-A
eFluor780
CD45 AF488
Live CD45+
FMO - No CD44
TN
TCM
TEM
CD62L BV786
CD44 PerCP-Cy5.5
CD4 BV510
CD8α AF647
CD8+CD4-
FMO - No Pentamer PE
Pentamer+ CD8+ TCM
Pentamer+ CD8+ TEM
Pentamer+ CD8+ T cells
Pentamer PE
Naive
ATRi QDx3 + RT

**Figure S10. Flow cytometry gating strategy to identify tumor antigen-specific $CD8^+$ T cells in the DLN.**

Any unstable portions of the run were gated out prior to analysis. After single cell gating (FSC-A vs. FSC-H) to remove doublets/clumps and scatter gating (FSC-A vs. SSC-A) to exclude debris and, live immune cells (efluor780$^{neg}$ and $CD45^+$) in the DLN were gated for further analysis. $CD8^+CD4^-$ cells were examined for AH1 Pentamer binding to identify tumor antigen-specific (Pentamer$^+$) $CD8^+$ T cells. In addition, $CD8^+CD4^-$ cells were examined for expression of CD62L and CD44 to identify naïve ($CD62L^{hi}CD44^{lo}$), central memory (TCM, $CD62L^{hi}CD44^{hi}$), and effector/effector memory (TEM, $CD62L^{lo}CD44^{hi}$) $CD8^+$ T cells. TCM and TEM were further examined for AH1 Pentamer binding to identify tumor antigen-specific $CD8^+$ TCM and tumor antigen-specific $CD8^+$ TEM. Fluorescence-minus-one controls are shown for empiric determination of CD44 and Pentamer gates. DLN from a naïve (no tumor) control mouse, included with each experiment, is shown to demonstrate low background Pentamer binding. DLN from an ATRi QDx3 plus RT-treated mouse is shown to demonstrate positive Pentamer binding.

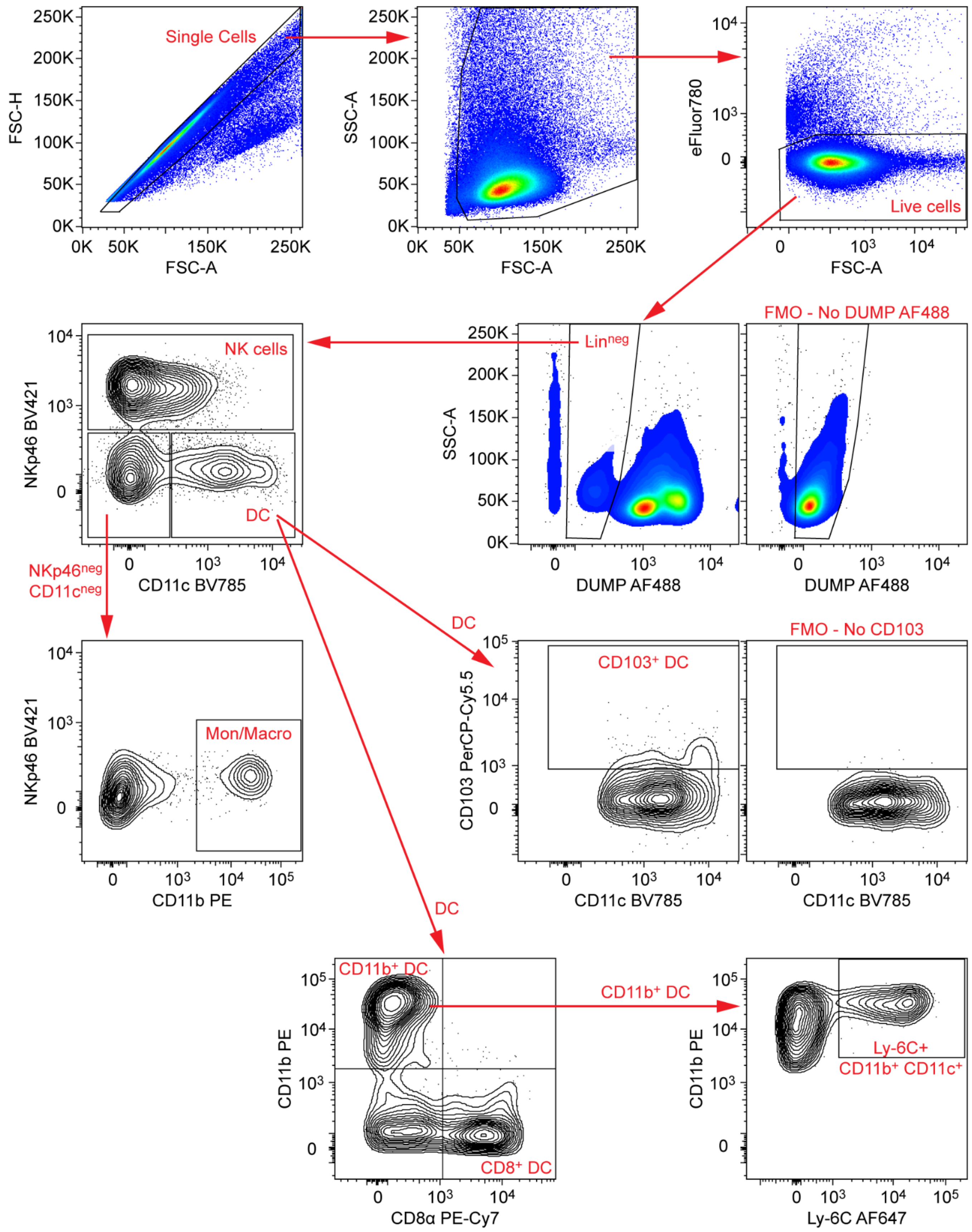

Single Cells
Live cells
FMO - No DUMP AF488
Lin^neg
NK cells
DC
NKp46^neg CD11c^neg
Mon/Macro
FMO - No CD103
CD103+ DC
CD11b+ DC
CD8+ DC
Ly-6C+ CD11b+ CD11c+
FSC-H
FSC-A
SSC-A
eFluor780
DUMP AF488
NKp46 BV421
CD11c BV785
CD11b PE
CD103 PerCP-Cy5.5
CD8α PE-Cy7
Ly-6C AF647

**Figure S11. Flow cytometry gating strategy to identify innate immune cells in the DLN.**

Any unstable portions of the run were gated out prior to analysis. After single cell gating (FSC-A vs. FSC-H) to remove doublets/clumps and scatter gating (FSC-A vs. SSC-A) to exclude debris, live cells (efluor780$^{neg}$) in the DLN were gated, and T and B cells (CD3$^{+}$ or CD19$^{+}$, DUMP channel) were excluded. The Lin$^{neg}$ (CD3$^{neg}$CD19$^{neg}$) cells were subset into NK cells (NKp46$^{+}$), dendritic cells (DC, CD11c$^{+}$ NKp46$^{neg}$) and CD11c$^{neg}$NKp46$^{neg}$. The CD11c$^{neg}$NKp46$^{neg}$ cells were further examined for CD11b expression to identify monocytes/macrophages (Mon/Macro). DC were further profiled for CD103, CD8α, or CD11b expression to identify CD103$^{+}$ DC, CD8$^{+}$ DC, or CD11b$^{+}$ DC, respectively. Ly-6C expression on CD11b$^{+}$ DC was also examined. FMO control plots are shown for DUMP and CD103.

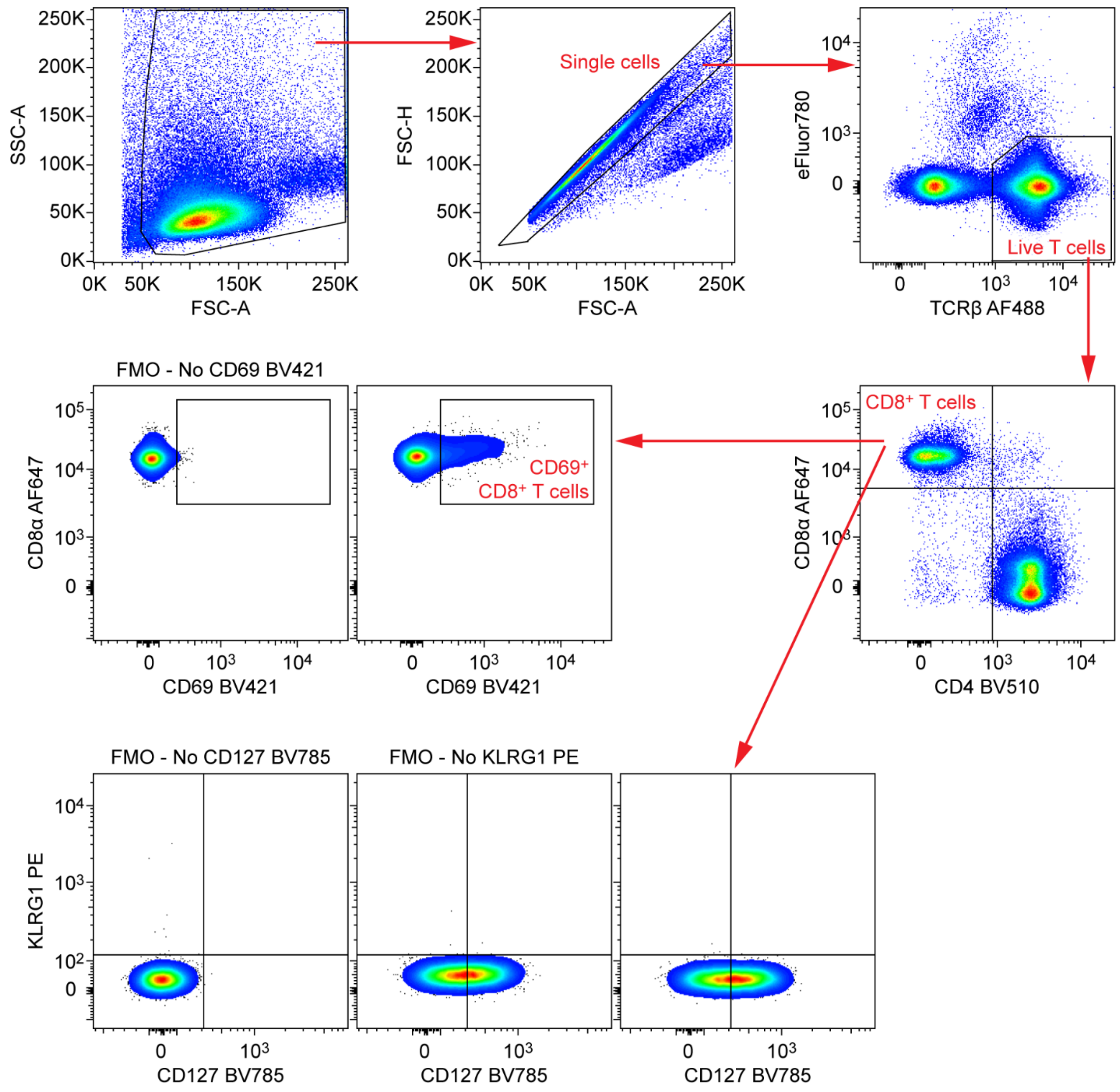


**Figure S12. Flow cytometry gating strategy to identify newly activated CD8$^+$ T cells in the DLN.**

Any unstable portions of the run were gated out prior to analysis. After scatter gating (FSC-A vs. SSC-A) to exclude debris and single cell gating (FSC-A vs. FSC-H) to remove doublets/clumps, live T cells (efluor780$^{neg}$ and TCR$^+$) in the DLN were gated and subset into CD8$^+$ T cells and CD4$^+$ T cells. CD8$^+$ T cells were further examined for expression of CD69 to identify newly activated CD8$^+$ T cells in the DLN. KLRG1 PE and CD127 BV785 were included in the staining panel, but CD8$^+$ T cells expressing these markers were not reported due to very low abundance of KLRG1$^+$ CD8$^+$ T cells and weak staining of CD127 on CD8$^+$ T cells. FMO control plots are shown for CD69, KLRG1, and CD127.
